# Auditory stimuli extend the temporal window of visual integration by modulating alpha-band oscillations

**DOI:** 10.1101/2024.01.31.578121

**Authors:** Mengting Xu, Biao Han, Qi Chen, Lu Shen

**Author notes:** Correspondence (Q.C.); (L.S.). These authors contributed equally (Mengting Xu, Biao Han).

## Abstract

In multisensory environments, signals from different modalities interact to shape perception, but the underlying temporal dynamics remain unclear. We investigated how a brief auditory stimulus affects the temporal integration of successive visual events using electroencephalography (EEG). Participants viewed two consecutive flashes, with or without an auditory beep accompanying the first flash, and reported whether they perceived one or two flashes. The beep shifted the psychometric fusion threshold to longer interstimulus intervals, indicating a wider temporal integration window. This behavioural shift was accompanied by a transient reduction in poststimulus instantaneous alpha frequency, and individual differences in alpha-cycle lengthening correlated with threshold prolongation. Auditory input also weakened the relationship between prestimulus alpha frequency and perceptual report while strengthening the corresponding relationship with prestimulus alpha phase. In a follow-up transcranial alternating current stimulation (tACS) experiment, the effects of lower- versus upper-alpha stimulation on fusion thresholds differed between the visual-only and audiovisual conditions. A computational model based on phase resetting and perceptual cycles qualitatively reproduced the main findings. Together, the results suggest that a brief sound can reorganize alpha-band timing and thereby widen the temporal window of visual integration.

## Introduction

In everyday life, we continuously receive information from multiple senses. Our brains face the challenge of organizing and integrating these signals into coherent perceptual experiences, and this becomes more complex due to the interactions between different sensory modalities (Cooke et al., 2019; Shams et al., 2002; Watkins et al., 2006). Numerous studies consistently highlight the noteworthy influence of sound signals on visual perception, particularly in the temporal domain. For example, the introduction of sound has been observed to influence the perceived duration of visual stimuli or rate of visual events (Gebhard and Mowbray, 1959; Shipley, 1964; Walker and Scott, 1981; Welch et al., 1986). The temporal relationship between visual and auditory stimuli either enhances or degrades visual temporal resolution (Fendrich and Corballis, 2001; Shimojo et al., 2001). These findings suggest that auditory stimuli strongly shape the temporal processing of the visual system. However, the neural mechanisms that underlie this cross-modal influence remain elusive.

The temporal integration window defines the range of separations over which distinct visual events can interact and be fused into a single percept (Samaha and Romei, 2024). Sensory uncertainty influences audiovisual simultaneity judgments, and stimulus reliability affects the temporal integration of successive visual inputs (Cary et al., 2024; Rangelov et al., 2021). In the double-flash paradigm, its width is indexed by the fusion-threshold ISI, defined as the separation at which one-flash and two-flash reports are equally likely. A higher threshold therefore indicates a wider integration window. Rhythmic neural oscillations provide a potential mechanism for defining this temporal window (Baumgarten et al., 2015; Cecere et al., 2015; Samaha and Postle, 2015). This notion gains support from research demonstrating that the phase and frequency of alpha oscillations shape the temporal integration process. Alpha oscillation phase influences visual signal detectability at the perceptual threshold (Busch et al., 2009; Busch and VanRullen, 2010; Mathewson et al., 2009), the perceived timing of stimuli (Chakravarthi and VanRullen, 2012; Milton and Pleydell-Pearce, 2016), and perceptual selection and temporal organization (Gulbinaite et al., 2017; Rohe et al., 2019; Ronconi et al., 2017). Emerging evidence links alpha frequency to temporal integration (Cecere et al., 2015; Cooke et al., 2019; Han et al., 2023; Samaha and Postle, 2015; Shen et al., 2019). Such relationships have been reported both within (Han et al., 2023; Samaha and Postle, 2015; Shen et al., 2019) and across different sensory modalities (Cecere et al., 2015; Cooke et al., 2019; Keil and Senkowski, 2017). However, alpha–perception relationships have not been observed consistently across tasks and analysis approaches (Buergers and Noppeney, 2022; Gray and Emmanouil, 2020; Ruzzoli et al., 2019). This highlights the need for further research to confirm the relationship between alpha oscillations and the temporal window within and across the senses.

In the context of audiovisual processing, the presentation of two visual flashes with an auditory stimulus leads to an increased likelihood of perceiving them as a single flash, inducing fusion illusions (Andersen et al., 2004). This observation suggests that cross-modal stimulation can alter the temporal window of visual processing. Neural studies have revealed significant activation changes in the primary visual cortex (V1) when visual stimuli are presented concurrently with incongruent auditory stimuli, which are closely linked to reports of visual illusions (Watkins et al., 2006). Research in animals and humans further shows that cross-modal inputs can modulate ongoing activity in sensory cortex (Kayser et al., 2008; Keil et al., 2014; Lakatos et al., 2007; Mercier et al., 2013). Together, these studies provide a plausible route through which sound could influence visual timing, although only some directly examined auditory-driven resetting in visual cortex. We therefore tested whether a cross-modal sound influences the temporal window of visual integration through changes in alpha oscillations.

To test this hypothesis, we conducted an experiment in which observers performed a visual flash discrimination task while their EEG data were recorded. We analysed the frequency and phase of alpha oscillations for two experimental conditions: one with only two visual flashes (F2) and the other with two visual flashes accompanied by an auditory beep (F2B1) (Figures 1A and B). Behaviourally, the auditory beep shifted the fusion threshold to longer ISIs, indicating an extended temporal integration window. This behavioural extension was accompanied by a transient poststimulus reduction in instantaneous alpha frequency. In the F2 condition, instantaneous alpha frequency predicted the perceptual outcome, whereas in the F2B1 condition it did not. Conversely, alpha phase predicted the perceptual outcome in F2B1 but not in F2. A follow-up transcranial alternating current stimulation (tACS) experiment characterised the condition-dependent effects of stimulation frequency. To examine a possible mechanism, we developed a sound-induced phase-resetting model. The model qualitatively reproduced the condition-dependent frequency and phase patterns, supporting phase resetting as a candidate account.

**Figure 1.**
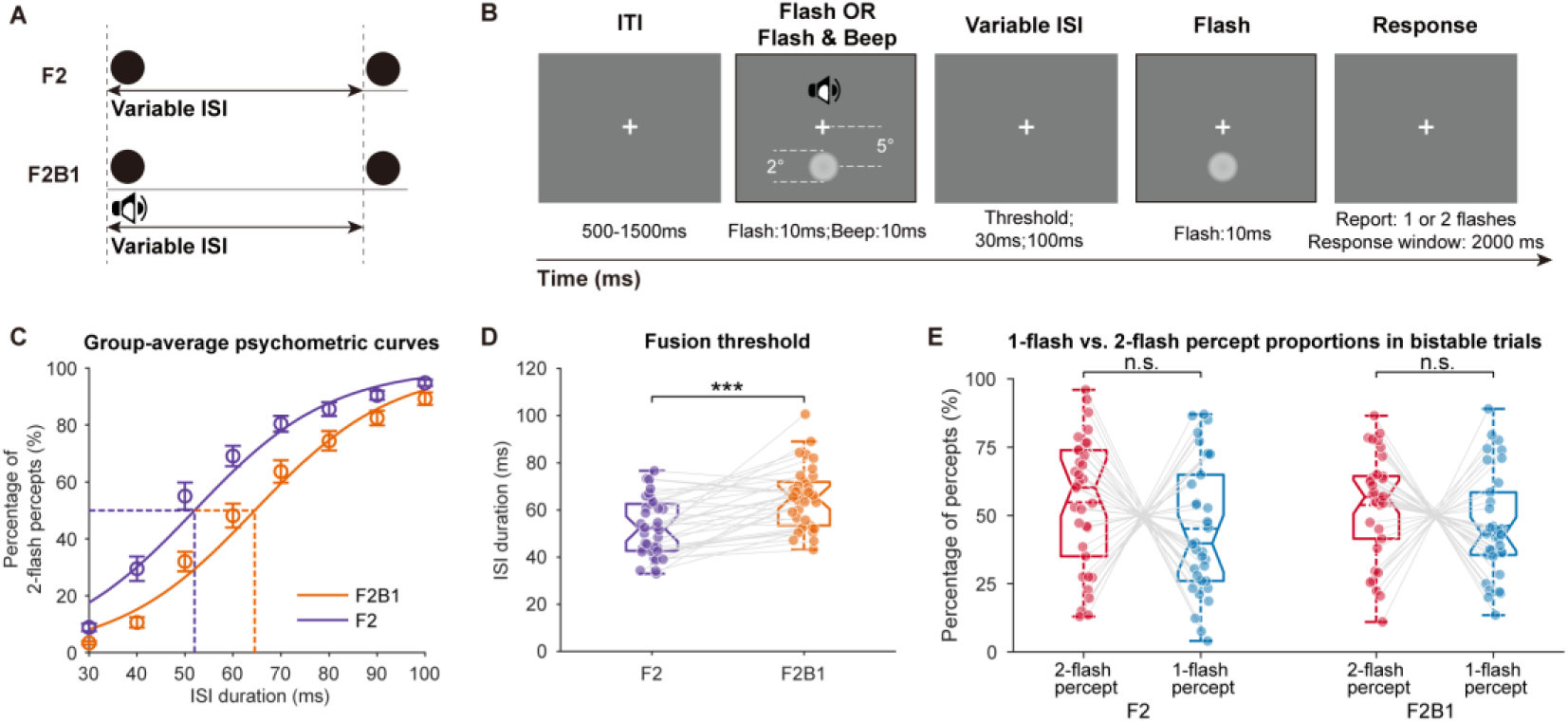
Paradigm and behavioural results. (A) Schematic of the two sensory conditions. In F2, two visual flashes were presented alone; in F2B1, the first flash was accompanied by an auditory beep. (B) Illustration of one trial sequence. Each trial began with a central fixation (500–1500 ms), followed by the first visual stimulus (10 ms; accompanied by the beep in F2B1), a variable ISI (30 ms, 100 ms, or the individually titrated fusion-threshold ISI), and the second visual stimulus (10 ms); participants then reported whether they perceived one or two flashes. The sound icon is illustrative and was not part of the display. (C) Group-average psychometric curves. Symbols show the mean percentage of two-flash reports at each ISI (mean ± *SEM*; error bars) for F2 (purple) and F2B1 (orange); continuous lines show binomial logistic fits to response counts pooled across participants, and the dashed lines mark the fitted 50% thresholds. (D) Fusion- threshold ISIs for each participant (dots) and condition; boxes show the median and interquartile range, whiskers the non-outlier range, dashed lines the mean, and grey lines the within-participant pairing between F2 and F2B1 (FDR-adjusted two-tailed paired test). (E) Percentage of valid bistable trials reported as one- flash (blue) and two-flash (red) percepts for F2 and F2B1; boxes show the median and interquartile range, whiskers the non-outlier range, dots individual participants, and grey lines connect the two report classes within each condition (FDR-adjusted two-tailed paired tests). \*\*\**p* < .001; n.s., not significant.

## Results

### Psychophysical Results

Prior to the main EEG experiment, participants underwent a psychophysical pretest to determine the fusion threshold for each condition. Psychometric curves were fitted to the proportion of two-flash reports at each of the eight ISIs for the F2 and F2B1 conditions, respectively (Figure 1C). The fusion threshold, representing the ISI at which one- and two-flash reports were equally likely, was derived for each condition and participant (Figure 1— figure supplement 1). This threshold provides an estimate of the temporal window for integration within the visual modality (F2) and across the audiovisual modality (F2B1). An extended integration window was operationalized as a rightward shift of this threshold: the two flashes could be separated by a longer ISI while remaining equally likely to be reported as one or two. A two-tailed paired *t*-test revealed that the fusion threshold was significantly shorter (*t*_(33)_ = −5.23, *p* < .001; Wilcoxon signed-rank *p* < .001; Figure 1D) in the F2 (52.68 ± 12.25 ms, mean ± *SD*) than in the F2B1 condition (65.19 ± 13.17 ms, mean ± *SD*), indicating a significant shift in the fusion threshold between the two conditions. Further analysis revealed that the auditory stimulus affected participants’ choices differently across ISIs (*F*_(7, 231)_ = 11.12, *p* < .001; Figure 1—figure supplement 2), with the strongest effect at intermediate ISIs, where the stimulus was most ambiguous. Together with the counterbalanced response mapping and within-participant design, these findings argue against a simple, condition-independent response bias as an explanation of the auditory effect.

In the main EEG experiment, each condition comprised three types of trials: the explicit short-ISI trials (30 ms, typically reported as one flash), the explicit long-ISI trials (100 ms, typically reported as two flashes), and the bistable trials with the fusion threshold ISI obtained from the psychophysical procedure. In the fixed-ISI trials, the beep shifted reports toward fusion: it increased accuracy for the expected one-flash report at 30 ms but decreased accuracy for the expected two-flash report at 100 ms (Figure 1—figure supplement 3A; see Supplementary Results). Regarding the bistable trials with the threshold ISI, the proportions of 2-flash and 1-flash percepts were comparable in both the F2 (54.8% ± 4.1% versus 45.2% ± 4.1% (mean ± *SEM*), *t*_(33)_ = 1.18, *p* = .248, two-tailed, *BF*_01_ = 2.89) and F2B1 conditions (53.8% ± 3.3% versus 46.2% ± 3.3%, *t*_(33)_ = 1.16, *p* = .255, two-tailed, *BF*_01_ = 2.94; Figure 1E), suggesting that participants experienced a similar level of perceptual ambiguity in the two conditions. Consistent with this, reaction times in the bistable trials did not differ between conditions (F2, 725.29 ± 19.19 ms; F2B1, 733.35 ± 19.39 ms; *t*_(33)_ = −0.95, *p* = .350, Cohen’s *d_z_* = −0.16, *BF*_01_ = 3.60; Wilcoxon signed-rank *p* = .174; Figure 1—figure supplement 4). Because the bistable ISI was titrated separately for each condition, the two conditions were equated on perceptual ambiguity by construction, so comparable decision times are expected.

### Auditory input extended the temporal window for visual integration by decreasing occipital alpha frequency

The mean participant-level alpha peak was 10.12 ± 0.24 Hz (mean ± *SEM*), obtained as the 8– 13 Hz maximum of each participant’s trial-averaged FFT spectrum. Alpha power showed a posterior scalp distribution during the peri-stimulus period (−600 to 600 ms relative to the first stimulus onset) across participants in both F2 and F2B1 conditions (Figure 2A). These results supported the selection of the 8–13-Hz frequency band and a posterior electrode region of interest (PO7, PO5, PO3, POz, PO4, PO6, PO8, O1, Oz, and O2) for subsequent analyses.

**Figure 2.**
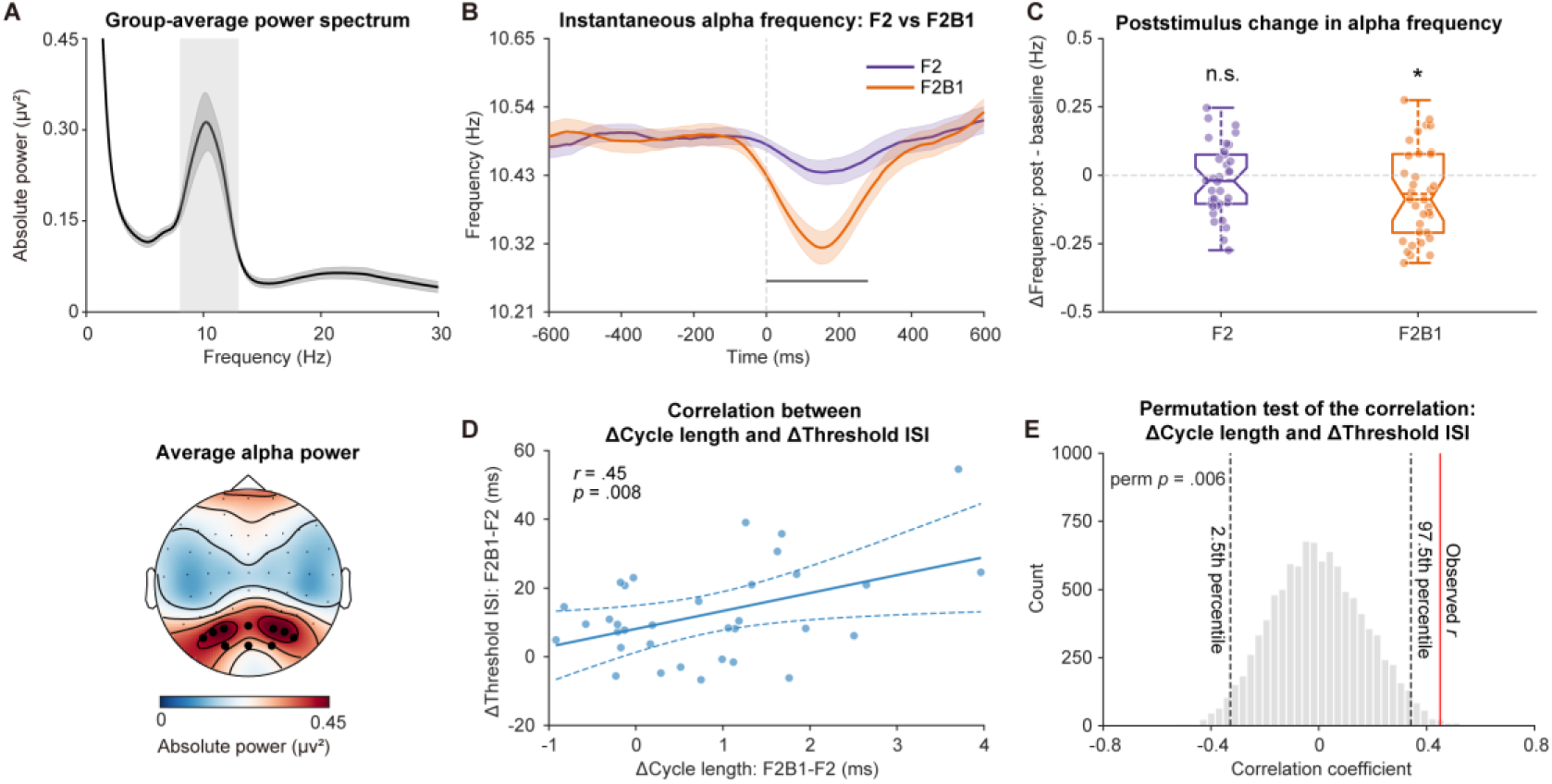
Auditory-induced poststimulus alpha-frequency slowing and fusion-threshold extension. (A) Group-average power spectrum from all bistable trials (−600 to 600 ms relative to the first flash onset) pooled across F2 and F2B1 (black), all electrodes, and participants; the topography below shows absolute alpha-band power (8–13 Hz) averaged over F2 and F2B1. Shaded areas show ±1 within-subjects *SEM*, the light grey rectangle marks the 8–13 Hz alpha band, and black dots mark the posterior channels used for the instantaneous-frequency analyses (PO7, PO5, PO3, POz, PO4, PO6, PO8, O1, Oz, and O2). (B) Time-resolved instantaneous alpha frequency (IAF) for F2 (purple) and F2B1 (orange) in the bistable trials, averaged over the posterior region of interest; shaded areas show ±1 within-subjects *SEM*, and the black horizontal bar marks the significant F2B1−F2 cluster (0–280 ms). (C) Poststimulus change in alpha frequency (0–600 ms minus the −600 to −10 ms baseline) for the bistable trials of each condition; boxes show the median and interquartile range, whiskers the non-outlier range, dashed lines the mean, and dots individual participants (FDR-adjusted one-tailed tests). (D) Between-subject Pearson correlation between the auditory-induced change in poststimulus alpha-cycle length (F2B1 − F2; calculated as 1,000/mean IAF over 0–280 ms in each condition before taking the difference) and the corresponding change in fusion- threshold ISI (F2B1 − F2); the solid line is the least-squares fit and the dashed curves show the 95% Working–Hotelling simultaneous confidence band for the fitted line (uncorrected correlation *p* value). (E) Null distribution of the correlation coefficient from 10,000 random re-pairings of the two measures; the red line marks the observed coefficient and the dashed black lines mark the 2.5th and 97.5th percentiles of the null distribution. \**p* < .05; n.s., not significant.

Having established that auditory input behaviourally extended the integration window, we next asked whether this extension was accompanied by a slowing of occipital alpha activity. If alpha frequency indexes the temporal integration window in the visual cortex, it should be lower in F2B1 than in F2. We therefore directly compared poststimulus instantaneous alpha frequency (IAF) between the two conditions, pooling 1-flash and 2-flash percepts while matching trial counts (see Materials and Methods). In bistable trials, poststimulus IAF was lower in F2B1 than in F2 from 0 to 280 ms after the auditory event. This difference was significant after two-tailed cluster-based correction (*p* = .006; Figure 2B). The reduction in instantaneous frequency indicates that alpha phase progressed more slowly in F2B1. We also examined the change in alpha frequency relative to baseline during bistable trials. Averaged over 0–600 ms, frequency decreased relative to the −600 to −10 ms baseline in F2B1 (Δ = −0.068 Hz, *t*_(33)_ = −2.39, FDR-corrected *p* = .023). The corresponding decrease in F2 did not reach significance (Δ = −0.019 Hz, *t*_(33)_ = −0.84, FDR-corrected *p* = .202; Figure 2C). The between-condition difference was also present in the explicit short- and long-ISI trials, in which the two conditions contained identical visual sequences (Figure 2—figure supplement 1A and B). Although alpha frequency also decreased relative to baseline in F2 at the long ISI (Figure 2—figure supplement 1C; see Supplementary Results), the F2B1−F2 contrast isolates the additional slowing associated with the beep.

A complementary FFT analysis provided convergent evidence. For each participant, we identified the frequency with maximal alpha power within 8–13 Hz during the 0–400-ms poststimulus interval. Peak frequency was lower in F2B1 (8.69 ± 0.18 Hz) than in F2 (9.40 ± 0.23 Hz), *t*_(33)_ = 3.49, *p* = .001, Cohen’s *d_z_* = 0.60. This difference was also supported by a Wilcoxon signed-rank test (*p* < .001; Figure 2—figure supplement 2A). Together, the time-resolved IAF and FFT analyses indicate that auditory input was associated with a transient slowing of poststimulus occipital alpha activity.

We next asked whether auditory-induced alpha slowing was related to the behavioural extension across participants. The increase in poststimulus alpha-cycle length (ΔCycle = F2B1 − F2; the inverse of mean IAF) correlated positively with the increase in fusion-threshold ISI (F2B1 − F2; Pearson *r* = .45, *p* = .008, two-tailed; bootstrap 95% CI [.08, .68]; Figure 2D). Thus, participants with greater auditory-induced alpha slowing showed a larger extension of the fusion threshold. This association was supported by several robustness checks. It remained significant in a 10,000-iteration permutation analysis (two-tailed *p* = .006; Figure 2E) and a bisquare robust regression (slope = 4.30, *p* = .033, two-tailed). Leave-one-participant-out correlations were also positive in every iteration (*r* = .29–.50) and significant in 33 of 34 iterations (Figure 2—figure supplement 3). Together, these results link auditory-induced poststimulus alpha slowing to the behavioural extension of visual temporal integration.

### Auditory input disrupted the predictive role of prestimulus alpha frequency

Prestimulus alpha frequency has been shown to influence the outcome of visual integration. We therefore asked whether its relationship with perceptual outcome was preserved when the first flash was accompanied by a beep. Within each condition, we compared time-resolved instantaneous alpha frequency (IAF) between bistable trials perceived as one versus two flashes. In F2, IAF was higher on 2-flash than on 1-flash trials during both the prestimulus period (−330 to −30 ms relative to the first stimulus onset, *p* = .013, two-tailed cluster-based correction) and the poststimulus period (130–540 ms, *p* = .003, two-tailed cluster-based correction; Figure 3A). Consistent with previous findings, lower alpha frequency was therefore associated with integrating the two stimuli into a single percept, as expected if slower alpha defines a broader temporal integration window (Samaha and Postle, 2015). By contrast, IAF did not differ between perceptual outcomes in F2B1 (Figure 3B). After removal of the aperiodic 1/f component, the frequency–percept difference in F2 remained similar in direction and magnitude to the primary result. The overall pattern was again consistent with a stronger frequency–percept relationship in F2 than in F2B1 (Figure 3—figure supplement 1A–C; see Supplementary Results).

**Figure 3.**
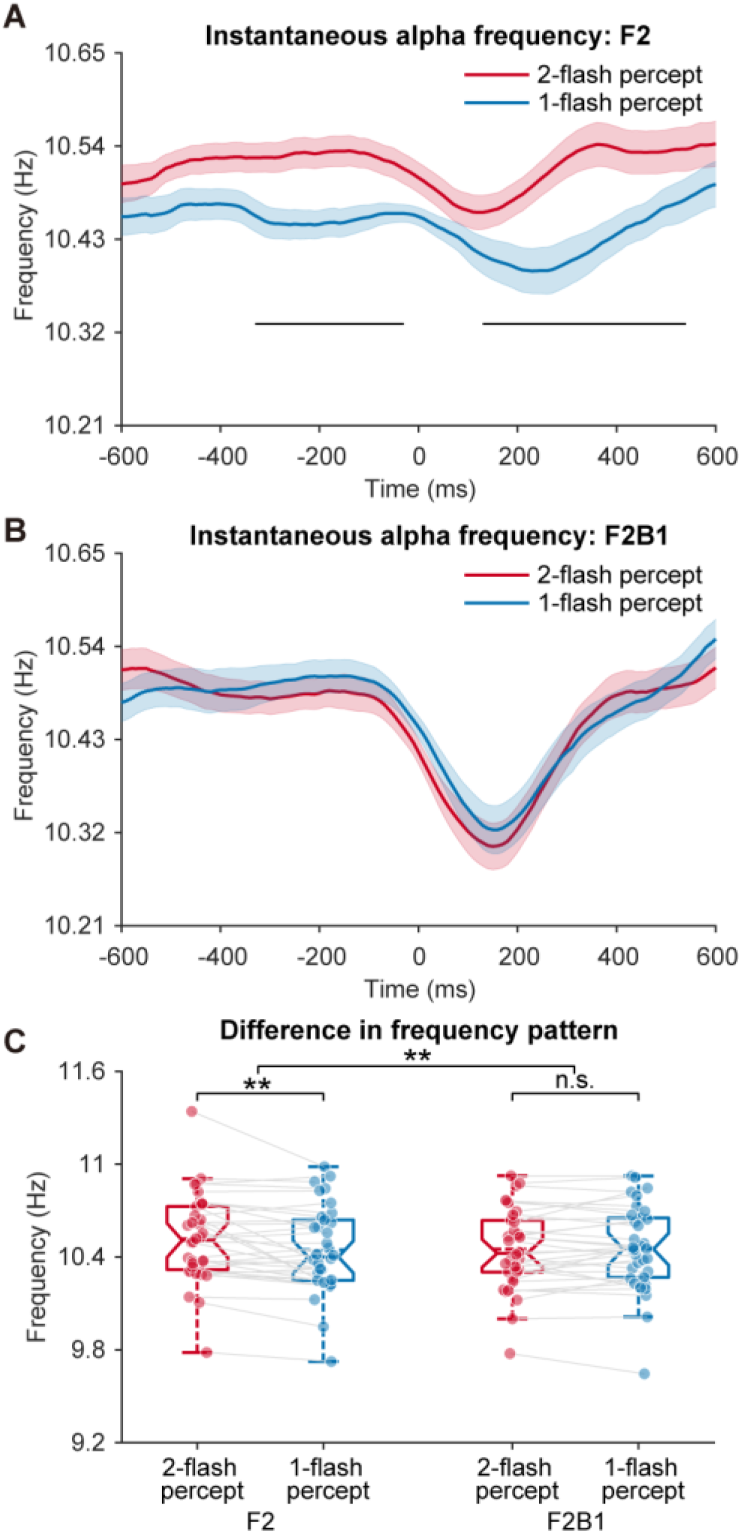
Relationship between instantaneous alpha frequency and perceptual report. (A) Time- resolved IAF over posterior channels for one-flash (blue) and two-flash (red) reports in F2; shaded areas show ±1 within-subjects *SEM* and black horizontal bars mark cluster-corrected significant periods. (B) The same analysis for F2B1. (C) Participant-level distributions and within-participant summaries of mean IAF for each percept and condition; boxes show the median and interquartile range, whiskers the non-outlier range, dashed lines the mean, dots individual participants, and grey lines the within-participant pairing between the two percepts (interaction: uncorrected repeated-measures ANOVA; within-condition comparisons: FDR-adjusted two-tailed paired tests). \*\**p* < .01; n.s., not significant.

To compare the frequency–percept relationships between conditions, we averaged IAF across the posterior electrodes and the full time window of interest (−600 to 600 ms). The resulting values were entered into a 2 × 2 repeated-measures ANOVA with perceptual outcome (1-flash versus 2-flash) and condition (F2 versus F2B1) as within-subject factors. The analysis revealed a main effect of perceptual outcome, *F*_(1, 33)_ = 4.29, *p* = .046, with higher IAF for 2-flash than for 1-flash percepts. The main effect of condition was not significant, *F*_(1, 33)_ = 3.40, *p* = .074, *BF*_10_ = 0.83. Critically, the interaction between perceptual outcome and condition was significant, *F*_(1, 33)_ = 9.86, *p* = .004. Follow-up comparisons showed that IAF was higher for 2- flash than for 1-flash percepts in F2, *t*_(33)_ = 3.11, uncorrected *p* = .004, FDR-corrected *p* = .008. In contrast, no difference between perceptual outcomes was detected in F2B1, *t*_(33)_ = −0.49, uncorrected and FDR-corrected *p* = .630, *BF*_01_ = 4.88 (Figure 3C). Thus, alpha frequency reliably distinguished perceptual outcomes in F2 but not in F2B1. Together with the time-resolved results, this interaction indicates that auditory input weakened the relationship between alpha frequency and perceptual outcome.

A complementary FFT analysis examined peak frequency during the peri-stimulus interval (−400 to 400 ms). The condition × percept interaction was significant, *F*_(1, 33)_ = 10.24, *p* = .003. In F2, peak frequency was higher for 2-flash than for 1-flash reports (*t*_(33)_ = 1.99, one-tailed uncorrected *p* = .027, FDR-adjusted *p* = .055; Wilcoxon signed-rank *p* = .009). In F2B1, peak frequency did not differ between perceptual outcomes (*t*_(33)_ = −1.31, one-tailed uncorrected *p* = .900; Wilcoxon signed-rank *p* = .488; Figure 2—figure supplement 2B). The significant interaction supports a stronger frequency–percept relationship in F2 than in F2B1, consistent with the instantaneous-frequency results.

To determine whether the IAF effects could be explained by differences in oscillatory power, we compared instantaneous alpha power between perceptual outcomes within the corresponding time window and electrodes of interest (Figure 3—figure supplement 2). Mean alpha power did not differ between 1-flash and 2-flash reports in either F2, *t*_(33)_ = −0.35, uncorrected and FDR-corrected *p* = .727, *BF*_01_ = 5.14, or F2B1, *t*_(33)_ = 1.04, uncorrected *p* = .308, FDR-corrected *p* = .616, *BF*_01_ = 3.31 (Figure 3—figure supplement 2C). Thus, the frequency effects reported above were unlikely to reflect concomitant differences in alpha amplitude.

### Auditory input enhanced the predictive role of prestimulus alpha phase

The spontaneous alpha-band phase has previously been shown to mirror the cyclic alterations in cortical excitability, thereby supporting the concept of perceptual cycles. We therefore asked whether prestimulus alpha phase predicts perceptual outcomes, using the phase opposition sum (POS), a measure of how much the phase distributions of two trial sets differ (VanRullen, 2016a). If the phase of spontaneous alpha oscillations before the first flash predicts the perceptual outcome, 1-flash trials should cluster around one phase angle and 2-flash trials around another, yielding a significant POS.

In the F2B1 condition, we observed a significant phase-opposition cluster between 1-flash and 2-flash trials, spanning −460 to 0 ms relative to the first flash onset and 5.34 to 20.89 Hz (*p* = .0002, one-tailed cluster-based correction; Figure 4A, upper panel). However, no significant phase effect emerged in the F2 condition (Figure 4A, lower panel). In line with the POS analysis, a Watson–Williams test showed that the preferred-phase directions differed between 1-flash and 2-flash percepts across participants in F2B1 (*F*_(1, 66)_ = 15.31, *p* < .001; Figure 4B, upper panel), whereas no such difference was found in F2 (*F*_(1, 66)_ = 3.30, *p* = .074; Figure 4B, lower panel).

**Figure 4.**
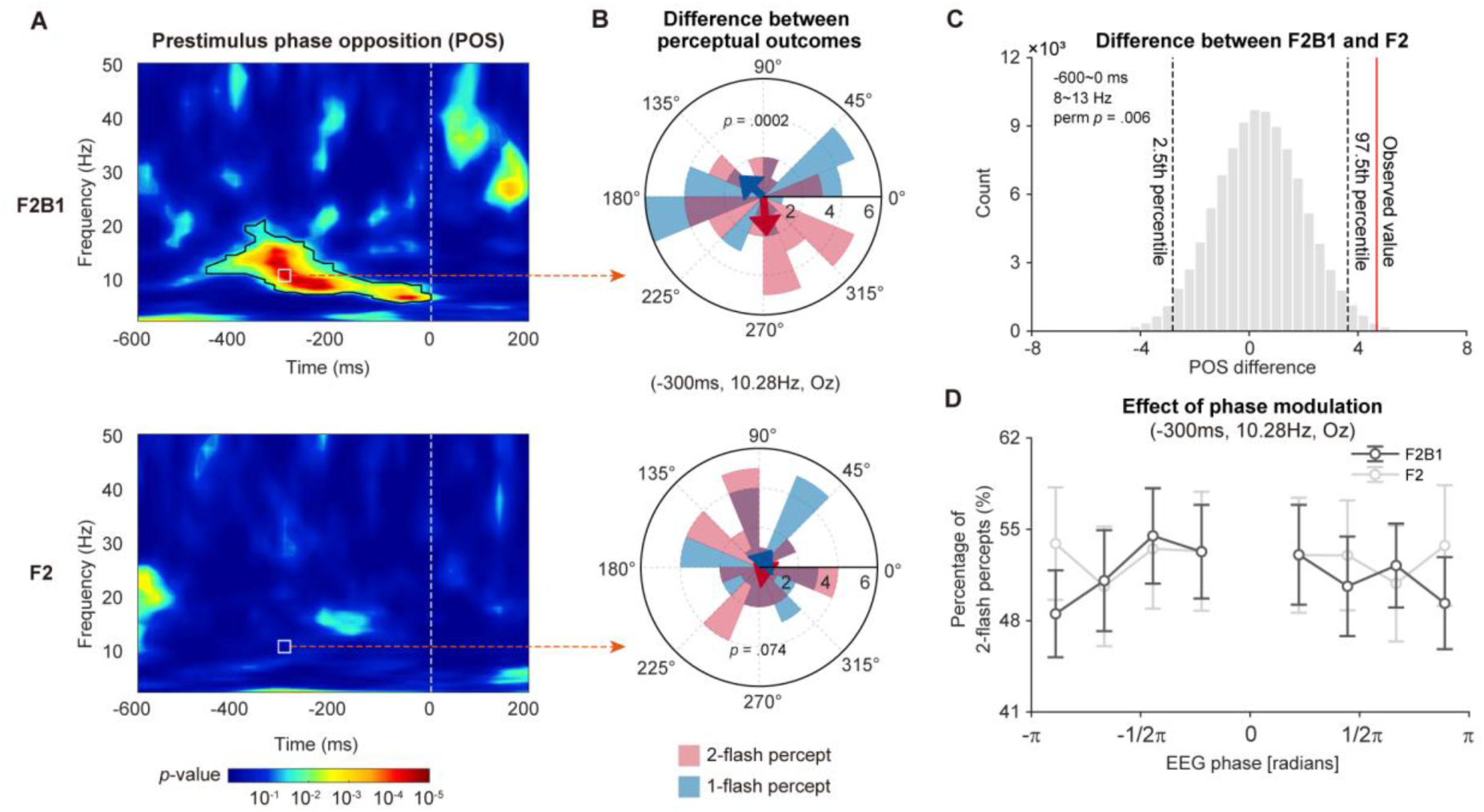
Prestimulus phase opposition between one-flash and two-flash reports. (A) Time–frequency maps of pointwise phase-opposition *p* values for F2B1 (top) and F2 (bottom); colour encodes −log10(*p*), with colour-bar tick labels showing *p* values from 10⁻¹ to 10⁻⁵ (colour bar at the bottom), the white dashed line marks stimulus onset, and black contours outline clusters surviving permutation-based correction. (B) Participant-level circular phase distributions at the illustrative maximum-effect point (−300 ms, 10.28 Hz, Oz) for one-flash (blue) and two-flash (red) reports in F2B1 (top) and F2 (bottom); arrows indicate group mean directions and the reported *p* values are from Watson–Williams tests of participant-level mean directions (uncorrected Watson–Williams tests). (C) Permutation null distribution of the F2B1 − F2 phase-opposition difference: the grey histogram shows the null distribution, the red line the observed value, and the dashed black lines the 2.5th and 97.5th percentiles; the y axis is count (×10³). (D) Phase-dependent two-flash report proportions (percentage) as a function of prestimulus phase at (−300 ms, 10.28 Hz, Oz) for F2B1 (dark) and F2 (light); symbols and error bars show the mean ± *SEM* across participants.

Furthermore, we evaluated whether the observed difference in prestimulus POS patterns between F2B1 and F2 was statistically significant using a permutation procedure. Specifically, we computed the summed POS difference between F2B1 and F2 in the frequency range of 8 to 13 Hz and time range from −600 to 0 ms relative to the first flash onset. Subsequently, we compared this summed POS difference value against a distribution of POS difference values expected by chance (see Materials and Methods; Figure 4C). This analysis confirmed a statistically significant distinction in POS patterns between F2B1 and F2 conditions (two-tailed permutation *p* = .006), indicating stronger phase modulation of perceptual outcomes in the F2B1 condition. To assess temporal smearing, we repeated the complete analysis using one-cycle wavelets. The one-cycle maps showed a similar prestimulus F2B1 alpha-band pattern without a comparable F2 pattern, reducing concern that the result was driven solely by wavelet support extending beyond stimulus onset (Figure 4—figure supplement 1).

We also examined whether the phase-opposition effect depended on prestimulus alpha power. POS was recomputed in high- and low-power subsets within each condition (Figure 4—figure supplement 2). In F2B1, the alpha-band cluster was significant in the high-power subset (cluster *p* = .004, −420 to −20 ms, 5.96–16.79 Hz) but not in the low-power subset; no alpha-band cluster appeared in either F2 subset, and the high-minus-low contrasts were nonsignificant.

Because these subset analyses were conducted separately within each condition, they are descriptive and do not formally test whether prestimulus alpha power modulates phase opposition differently in F2B1 and F2.

We further assessed the relationship between prestimulus phase and perceptual reports by sorting single trials according to phase at the optimal time–frequency point (−300 ms, 10.28 Hz, Oz) and calculating the proportion of 2-flash reports in each of nine phase bins (Figure 4D). For each participant, phase angles were shifted so that the bin with the highest proportion of 2-flash reports was centred at zero. Because this alignment necessarily makes the 2-flash proportion maximal at zero, the zero-phase bin was excluded from further analyses. If prestimulus phase modulates perceptual outcome, the proportion of 2-flash reports should decline as phase moves away from the optimal angle. We therefore tested the quadratic main effect of phase bin (eight levels after exclusion) and its interaction with condition (F2B1 versus F2). The condition × phase-bin interaction was significant (*F*_(1, 33)_ = 7.87, *p* = .008), indicating that quadratic phase modulation differed between conditions. The quadratic main effect of phase bin was not significant (*F*_(1, 33)_ = 3.33, *p* = .077). Follow-up tests showed a significant phase-modulation effect in F2B1, *F*_(1, 33)_ = 9.34, *p* = .004, but not in F2, *F*_(1, 33)_ = 0.04, *p* = .837 (Figure 4D).

A complementary non-aligned analysis quantified phase-dependent perceptual modulation from all artifact-free bistable trials, without assuming a common preferred phase. Each participant’s phase-response profile was summarized by its Kullback–Leibler (KL) divergence from a uniform profile and standardized against a participant-specific response-label permutation distribution. The mean standardized KL score was significantly above zero in F2B1 (mean *z* = 0.60, *t*_(33)_ = 2.32, one-tailed *p* = .013, FDR-corrected *p* = .029, one-tailed permutation *p* = .009) but not in F2 (mean *z* = −0.16, *t*_(33)_ = −0.91, one-tailed *p* = .816, FDR-corrected *p* = .816, one-tailed permutation *p* = .819; Figure 4—figure supplement 3A and B). The paired comparison confirmed stronger modulation in F2B1 (mean difference = 0.75, *t*_(33)_ = 2.46, two-tailed uncorrected *p* = .019, FDR-corrected *p* = .029; Figure 4—figure supplement 3C), and the permutation test on this difference corroborated the result (two-tailed permutation *p* = .014; Figure 4—figure supplement 3D).

### tACS-modulated alpha frequency causally influences the temporal integration window specifically in the F2 condition

Because EEG evidence is correlational, we conducted a follow-up experiment in a new group of participants using transcranial alternating current stimulation (tACS), which allowed us to modulate alpha frequency and test its causal role in perception.

During the experiment, continuous tACS was applied to the posterior brain region (Figure 5— figure supplement 1A) while participants performed the same task as in the psychophysical pretest, which required reporting their perception of two flashes across eight ISIs under the F2 and F2B1 conditions. Each participant completed three tACS sessions in a randomised order: low-alpha (8 Hz), high-alpha (13 Hz), and a sham condition with no active stimulation. Psychometric curves were fitted for each tACS session using the proportion of two-flash perceptions across eight ISIs, separately for F2 and F2B1 conditions. From these fitted curves, we calculated the fusion threshold ISI, the ISI at which participants perceived two flashes in 50% of trials, as an estimate of the temporal integration window (Figure 5A and B). Participant-level fitted curves and threshold estimates are shown in Figure 5—figure supplements 2 and 3.

**Figure 5.**
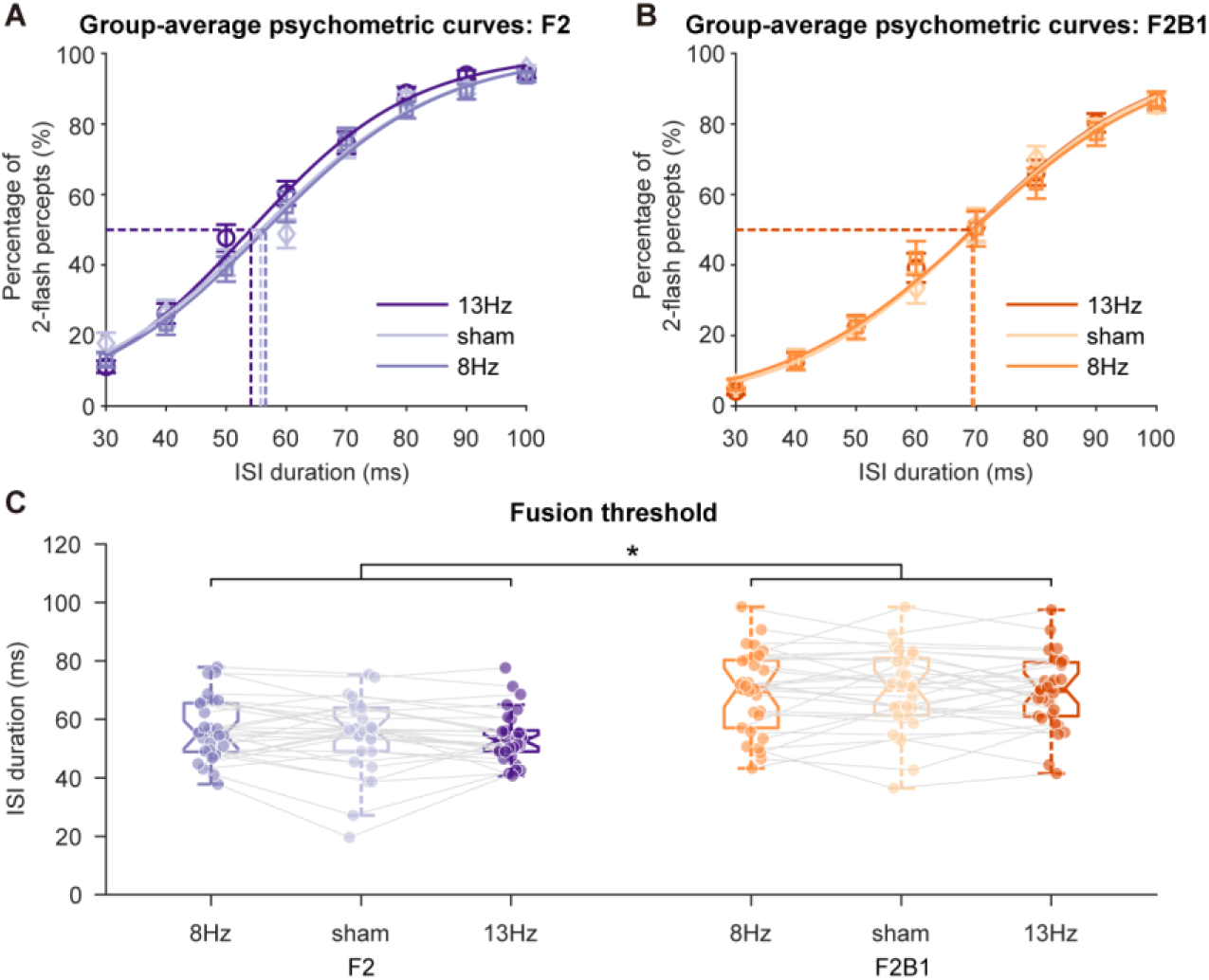
Results of the tACS experiment. (A) Group psychometric curves under 13 Hz (dark), sham (light), and 8 Hz (medium) stimulation in F2; symbols show the mean percentage of two-flash reports at each ISI and error bars ±1 *SEM*. (B) The same curves for F2B1. (C) Participant-level fusion thresholds across stimulation sessions (8 Hz, sham, 13 Hz) and sensory conditions (F2, F2B1); boxes show the median and interquartile range, whiskers the non-outlier range, dashed lines the mean, and dots individual participants; grey lines connect the sessions within each condition (8 versus 13 Hz × sensory condition interaction: repeated-measures ANOVA; sham shown for reference). Error bars represent ±1 *SEM* where applicable. \**p* < .05.

A 2 (stimulation frequency: 8 Hz vs. 13 Hz) × 2 (stimulus condition: F2 vs. F2B1) repeated-measures ANOVA on fusion thresholds revealed a significant main effect of stimulus condition, *F*_(1, 29)_ = 38.01, *p* < .001, consistent with the psychophysical pretest, showing that auditory input prolonged the temporal window for visual integration. The stimulation-frequency × sensory- condition interaction was significant (*F*_(1, 29)_ = 5.41, *p* = .027; Figure 5C), indicating that the effect of stimulation frequency on fusion thresholds differed between conditions. Because the difference scores were not normally distributed (Shapiro–Wilk *p* = .008), we confirmed the interaction with a two-tailed Wilcoxon signed-rank test on the (13 Hz − 8 Hz) difference between the two conditions, *p* = .043, and with a bootstrap 95% CI that excluded zero, [−5.56, −0.70] ms. The main effect of stimulation frequency was not significant (*F*_(1, 29)_ = 1.25, *p*= .272). In the F2 condition, the fusion threshold at 13 Hz was significantly lower than at 8 Hz (53.79 ± 1.62 ms vs. 56.45 ± 1.93 ms; mean difference −2.66 ms), *t*_(29)_ = −2.09, one-tailed *p* = .023, FDR-corrected *p* = .045, Cohen’s *d_z_* = −0.38, two-sided *BF*_10_ = 1.29; a one-tailed Wilcoxon signed-rank test, *p* = .034, and a bootstrap 95% CI that excluded zero, [−5.37, −0.40] ms, supported the same conclusion: higher alpha frequencies shortened the temporal window for perceptual integration. In contrast, in the F2B1 condition, the two stimulation frequencies produced indistinguishable thresholds (69.64 ± 2.30 ms vs. 69.40 ± 2.59 ms; mean difference 0.24 ms), *t*_(29)_ = 0.20, one-tailed *p* = .578, FDR-corrected *p* = .578, Cohen’s *d_z_* = 0.04, two-sided *BF*_01_ = 5.05. A sham session was also collected and is plotted for reference (F2, 54.88 ± 2.26 ms; F2B1, 69.87 ± 2.55 ms); because it carries no frequency prediction, it was not entered into the statistical model.

### An auditory-induced phase-resetting model reproduces the frequency and phase effects

What could underlie the decrease in poststimulus alpha frequency in F2B1, and could the same mechanism explain the altered predictive roles of alpha frequency and phase? Cross-modal inputs can reset ongoing oscillations in sensory cortex (Lakatos et al., 2009, 2007), and intracranial studies show that such phase reorganization contributes to multisensory processing (Mercier et al., 2015, 2013). Such a reset shifts the timing of the ongoing oscillation, which could in turn alter its apparent speed. In the model, the auditory event delays the ongoing phase trajectory; because instantaneous frequency is the local temporal derivative of phase, this delay temporarily lengthens the current cycle and appears as a brief reduction in instantaneous frequency shortly after the auditory event. The slowing is time-limited, recovering as the trajectory continues, and is distinct from a sustained change in trait-like individual alpha frequency.

To test this hypothesis, we introduced a phase-resetting model for examining perceptual processing (Figure 6A). Built on the perceptual-cycle framework (VanRullen, 2016b), the model uses simulations to replicate our key observations. Within this framework, frequency and phase jointly shape perceptual outcomes. Frequency determines whether the two flashes fall within the same or different alpha cycle, thereby biasing the potential perceptual outcome: higher frequencies favour the perception of two distinct flashes, while lower frequencies tend to merge them into a single event. Phase determines how reliably that outcome is reported. Our model differentiates percepts by assessing whether two flash stimuli fall within a single alpha cycle. Phases near the peak enhance processing efficiency, making 1-flash and 2-flash percepts clearer; phases near the trough reduce efficiency and increase perceptual ambiguity (Figure 6B). In F2B1, the auditory event delays the ongoing phase trajectory, and the model is intended to test whether this additional auditory perturbation can provide a simplified mechanistic account of the observed condition differences.

**Figure 6.**
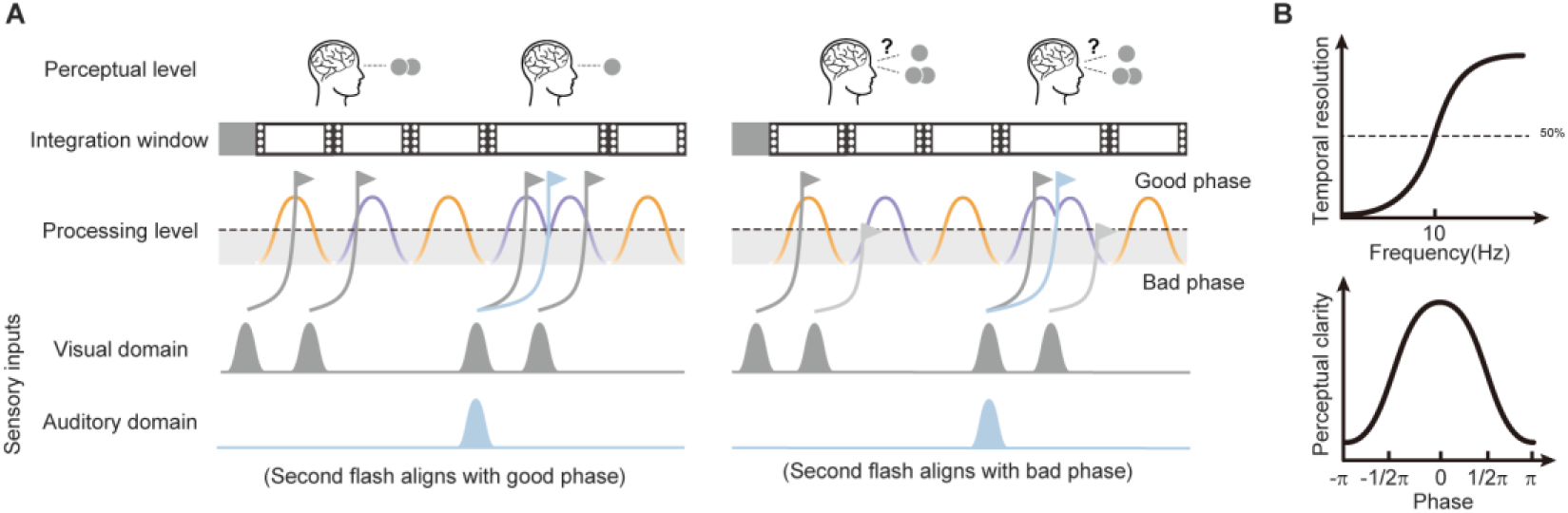
Hypothesized phase-resetting model. (A) Mechanism diagram: alpha frequency determines whether the two flashes occupy the same or different cycles and thus biases the latent one-flash versus two-flash outcome; the auditory event delays the ongoing phase trajectory, transiently lengthening the current cycle and increasing the ISI over which the flashes can be fused. The left column shows the case in which the second flash aligns with a good phase (high-excitability phase), yielding clear one-flash or two-flash percepts; the right column shows the bad-phase case, in which the second flash falls at a low-excitability phase and the report becomes less reliable. (B) Conceptual functions relating the two model roles: the upper plot shows temporal resolution as an increasing function of frequency (two flashes fall in the same or different cycles depending on the cycle length), and the lower plot shows perceptual clarity as a function of phase, with the highest clarity at zero phase and lower clarity toward ±π. Frequency therefore concerns perceptual outcome, whereas phase concerns the fidelity with which that outcome is reported.

Our data are consistent with this account. Alpha inter-trial coherence (ITC) was higher in F2B1 than in F2 from 40 to 220 ms (cluster *p* = .005, two-tailed; Figure 7A), and mean ITC over the 0–600-ms poststimulus interval was also higher in F2B1, *t*_(33)_ = 3.67, *p* < .001 (Figure 7B). In bistable trials, the F2B1−F2 instantaneous-frequency difference lasted from 0 to 280 ms (cluster *p* = .006, two-tailed), spanning more than two cycles at the approximately 10-Hz group alpha peak. Because an increase in ITC following a stimulus may reflect a stimulus-evoked response, we also examined changes in alpha power after stimulus onset. An additive evoked response would increase both ITC and power, whereas pure phase resetting should increase ITC without increasing power (Makeig et al., 2004; Shah et al., 2004). No alpha-power cluster survived correction (Figure 7C), and the between-condition differences in mean alpha power and ITC over the 0–600-ms interval were not significantly correlated, *r* = .12, *p* = .507 (Figure 7D). Therefore, although scalp EEG cannot fully distinguish phase resetting from other stimulus-evoked phase-locking mechanisms, the increase in alpha ITC is consistent with a transient reorganization of phase rather than a purely additive response.

**Figure 7.**
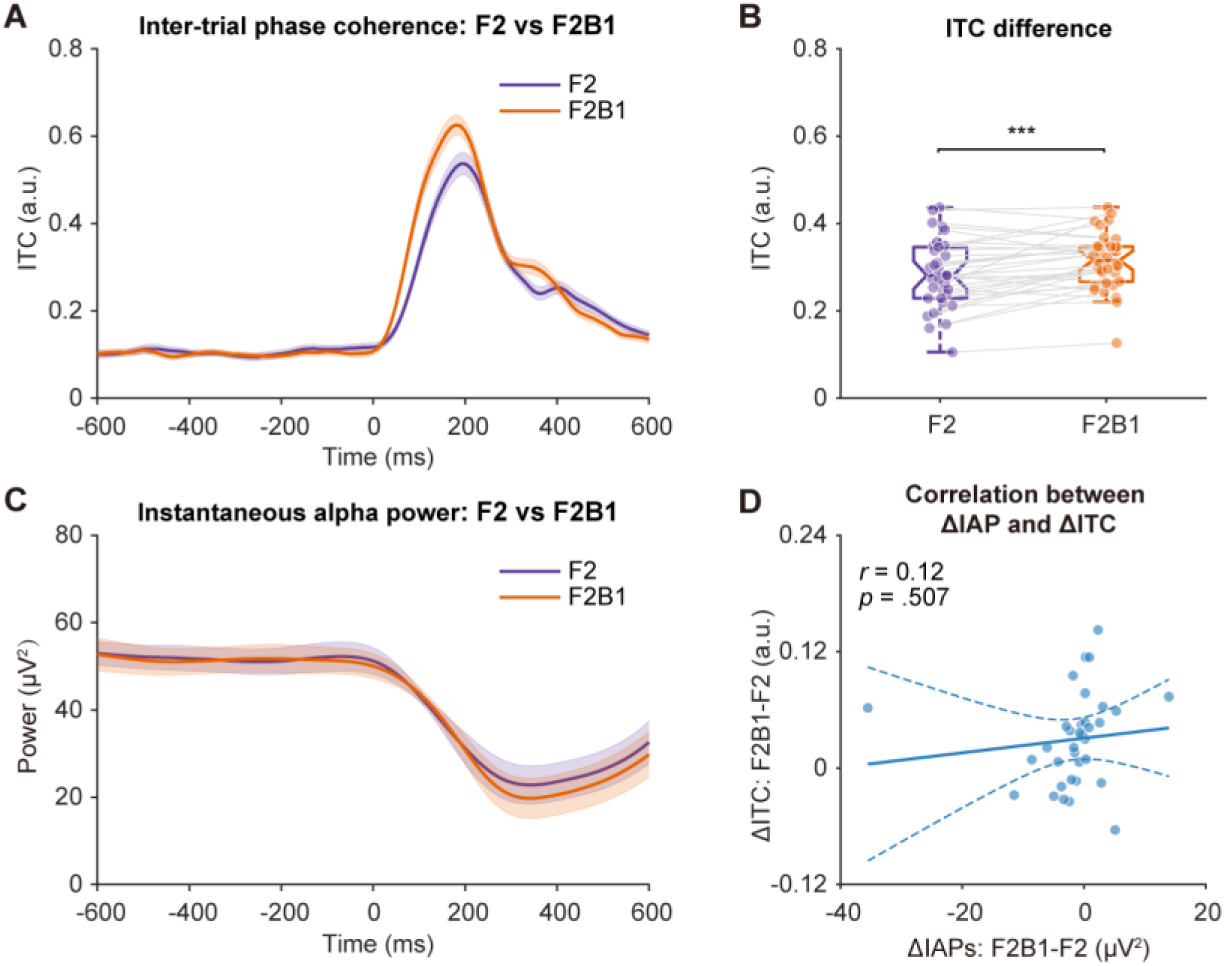
Poststimulus inter-trial phase coherence (ITC) and alpha-power control. (A) Time-resolved ITC for F2 (purple) and F2B1 (orange) in the bistable trials at the condition-specific fusion-threshold ISI; shaded areas show ±1 within-subjects *SEM*. (B) Mean ITC over 0–600 ms for F2 and F2B1; boxes show the median and interquartile range, whiskers the non-outlier range, dots individual participants, and grey lines the within-participant pairing (FDR-adjusted paired test). (C) Time-resolved instantaneous alpha power for F2 and F2B1; shaded areas show ±1 within-subjects *SEM*; no between-condition alpha-power cluster survived correction. (D) Between-subject correlation between the cross-condition ITC difference and the power difference (ΔIAP: F2B1 − F2; ΔITC: F2B1 − F2); the solid line is the least-squares fit, and dashed lines show its 95% confidence band. \*\*\**p* < .001.

Using the model workflow shown in Figure 8—figure supplement 1, we simulated 1,000 trials per condition. The simulated behavioural data reproduced the key pattern in the real data, with a prolonged integration window in F2B1 relative to F2 (Figure 8A). Applying the same analysis pipeline to the simulated oscillations reproduced the lower alpha frequency in F2B1; the modelled decrease was centred near stimulus onset and recovered within approximately 200 ms (Figure 8B; Figure 8—figure supplement 2). The frequency-percept separation was likewise reproduced: clear in F2 and smaller in F2B1 (Figure 8C). Sensitivity analyses varying the oscillator frequency and the phase delay supported the parameter choices made in the main simulation (Figure 8—figure supplements 3 and 4; see Supplementary Results).

**Figure 8.**
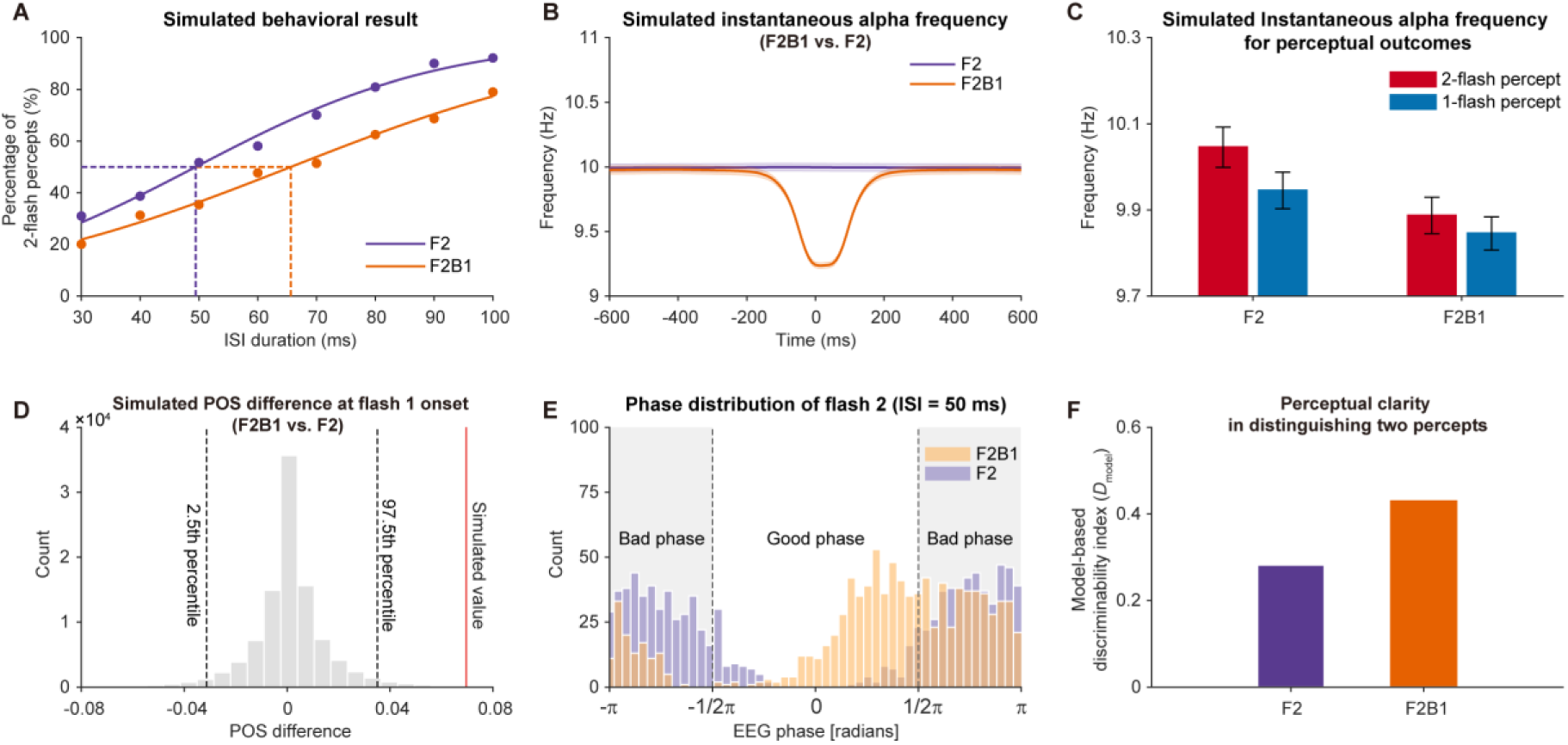
Simulation results. (A) Simulated psychometric curves and fusion thresholds for F2 (purple) and F2B1 (orange); symbols show the simulated percentage of two-flash reports at each ISI, continuous lines the logistic fits, and dashed lines the fitted 50% thresholds. (B) Time-resolved simulated IAF for F2 and F2B1; shaded areas show ±1 *SEM* across simulated trials, and the modelled decrease is centred near stimulus onset and recovers within approximately 200 ms. (C) Simulated IAF for two-flash (red) and one-flash (blue) perceptual outcomes in F2 and F2B1; bars show means and error bars ±1 *SEM* across simulated trials. (D) Simulated POS difference at flash-1 onset (F2B1 vs. F2): the grey histogram shows the null distribution from 100,000 permutations, the red line the simulated value, and the dashed black lines the 2.5th and 97.5th percentiles; the y axis is permutation count (×10⁴). (E) Phase distribution of the second flash at an ISI of 50 ms for F2 (purple) and F2B1 (orange); dashed lines mark the good-phase boundaries (±π/2). (F) Model-based discriminability index (*D*_model_), quantifying how closely the simulated reports corresponded to the model-defined latent one-flash and two-flash states; bars show the resulting *D*_model_ for F2 and F2B1.

The time-resolved phase analysis reproduced the condition-dependent phase separation observed in the empirical POS results. Phase opposition emerged around stimulus onset in both conditions and was stronger in F2B1 than in F2 (Figure 8D and Figure 8—figure supplement 5). The timing of the simulated effect differed from that of the empirical effect because poststimulus phase estimates in EEG may be obscured by stimulus-evoked phase locking. The empirical POS analysis therefore focused on the prestimulus period. By contrast, the model contained no stimulus-evoked response, so its poststimulus phase estimates directly reflected the ongoing oscillation and the imposed auditory phase delay. The simulated phase opposition was consequently locked to stimulus onset and was not expected to reproduce the precise prestimulus timing of the empirical F2B1 effect.

At the 50-ms ISI, the phase distributions of the second flash also differed in relation to the modelled good- and bad-phase regions (Figure 8E). This pattern was consistent with phase modulation being stronger in F2B1 but still present in F2. Finally, we quantified how accurately the simulated reports reflected the model-defined latent one-flash and two-flash states using the model-based discriminability index, *D_model_* (Equation 11). *D_model_* was higher in F2B1 than in F2, indicating a closer correspondence between the latent cycle-defined state and the simulated report in F2B1 (Figure 8F).

## Discussion

We examined how auditory stimuli influence the temporal window of visual integration. Auditory input prolonged the fusion threshold, decreased poststimulus alpha frequency in a way that correlated with the extended window, and engaged prestimulus alpha frequency and phase differently in unimodal versus cross-modal conditions. A computational model based on the phase-resetting hypothesis and perceptual-cycle theory (VanRullen, 2016b) qualitatively reproduced these key behavioural and neural findings. Together with the tACS interaction, these findings suggest that alpha frequency and phase make context-dependent contributions to visual temporal integration.

Our findings reveal that concurrent auditory input significantly extends the temporal window of visual integration compared to the no-sound condition (Figure 1C). This is consistent with earlier reports that sound alters the perceived timing of visual events (Gebhard and Mowbray, 1959; Shipley, 1964; Walker and Scott, 1981; Welch et al., 1986). This prolonged temporal window is positively correlated with the decrease in poststimulus alpha frequency induced by auditory stimuli. These findings contribute to the enduring hypothesis associating the human alpha rhythm with temporal windows, positing that higher frequencies correspond to shorter windows and heightened temporal resolution (Cecere et al., 2015; Cooke et al., 2019; Keil and Senkowski, 2017; Samaha and Postle, 2015). A lengthened integration window, as indicated by a longer alpha cycle, implies that two successive visual flashes are more prone to falling within a single cycle. Consequently, the visual system requires an extended duration to effectively differentiate between the two flashes.

In the present study, prestimulus alpha frequency predicted subsequent reports in the visual-only F2 condition, but this relationship was not reliable in F2B1. This condition difference should not be taken to mean that prestimulus oscillatory timing is unimportant for audiovisual fusion. Rather, it raises the possibility that the auditory event may change the frequency range through which preparatory temporal dynamics influence perception. Consistent with this possibility, Noguchi (2022) reported that lower prestimulus beta frequency was associated with greater susceptibility to audiovisual fusion. The present F2 result implicates alpha-band timing in purely visual double-flash perception; together, these findings raise the possibility that, when the beep contributes to fusion, the relevant timing information may shift to the beta band. Whether beta indeed takes over this role is a question for future work.

This raises a central question: how does auditory stimulation produce a decrease in alpha frequency? Previous work has shown that auditory stimuli can modulate activity in the primary visual cortex (Watkins et al., 2007, 2006) and that cross-modal inputs can reorganize ongoing sensory-cortical oscillations (Lakatos et al., 2009; Mercier et al., 2013; Romei et al., 2012), providing a plausible route by which the beep could alter the alpha rhythm. Consistent with this account, the accompanying increase in inter-trial phase coherence, without a corresponding power change, points to a reorganization of phase rather than an additive evoked response (Figure 7A and C). A phase reset of this kind is transient: it briefly slows the instantaneous frequency and recovers within a few cycles, which accounts for the time-limited decrease in the data without requiring a lasting change in alpha frequency. Because the F2 and F2B1 trials shared identical visual sequences, visual contributions cancel in the F2B1−F2 difference, leaving the auditory contribution introduced by the beep, together with any audiovisual interaction, as the source of the difference. Accordingly, the temporal extent of the F2B1−F2 difference is best interpreted as the interval over which the two conditions are statistically distinguishable; its relation to the duration of phase reorganization remains uncertain.

Moreover, our findings indicate that the predictive role of prestimulus alpha frequency depends on sensory condition. Consistent with previous studies (Han et al., 2023; Samaha and Postle, 2015; Shen et al., 2019; Wutz et al., 2018), higher alpha frequencies in the F2 condition were associated with a shorter integration window and more rapid sampling, enabling accurate discrimination of temporally close stimuli. In contrast, lower frequencies corresponded to a longer integration window and slower sampling, leading to fusion illusions (Figure 3A). This relationship was weaker and less reliable in F2B1, where auditory input may have reduced the predictive contribution of alpha frequency, consistent with recent null effects in an audiovisual context (Buergers and Noppeney, 2022). Phase showed the opposite pattern, with a stronger predictive role in F2B1 than in F2. The tACS results were consistent with this condition-dependent dissociation: stimulation frequency altered the fusion threshold in the visual-only condition, whereas a comparable modulation was not evident in the audiovisual condition under the present design. Together, the EEG and tACS results support a condition-dependent role for alpha frequency, with a stronger frequency effect in F2, while the EEG phase results show the opposite pattern. These findings do not exclude a contribution of alpha frequency to audiovisual temporal integration.

Prior research has shown that phase encodes rapidly changing stimuli effectively (Schyns et al., 2011), though temporal precision is often compromised by stimulus-evoked neural responses that dominate phase estimates (Harris, 2023; VanRullen, 2016a). As a result, prestimulus phase has become a valuable proxy for predicting perception, although its effectiveness has been debated. While some studies consistently support the predictive role of prestimulus phase (Busch et al., 2009; Gruber et al., 2014; Mathewson et al., 2009; Milton and Pleydell-Pearce, 2016), others report null effects (Benwell et al., 2022, 2017; Keitel et al., 2022; Ruzzoli et al., 2019; van Diepen et al., 2015). Within the model, frequency determines how much of an alpha cycle separates the two flashes, whereas phase influences how the resulting cycle-based state is expressed in the simulated report. Auditory-induced phase delay offers one account of why phase was more informative in F2B1; the simulated phase analysis likewise showed phase opposition in both conditions, stronger in F2B1 (Figure 8D).

The phase and frequency effects were not accompanied by differences in oscillatory amplitude (Figure 3—figure supplement 2 and Figure 7C), so they cannot be attributed to amplitude variations between conditions or percepts. Related work has linked several features of alpha activity, including phase and power, to perceptual processing (Jensen and Mazaheri, 2010; Mathewson et al., 2011, 2009; van Dijk et al., 2008; Williams et al., 2024). A separate question is whether the strength of the oscillation, as indexed by power, shapes how clearly these effects are expressed. Phase-related modulation was most apparent when the alpha oscillation was strongly expressed, in line with proposals that alpha power indexes the strength of the underlying rhythm and, with it, the reliability of phase information (Jensen and Mazaheri, 2010; Mathewson et al., 2011; van Dijk et al., 2008). Although the direct high- versus low-power comparison was not significant, this pattern is consistent with the idea that oscillatory strength modulates the expression of phase-based processing, a question that remains open for more targeted tests.

While there is substantial evidence supporting the role of alpha oscillation frequency in characterising the temporal window for integration (Han et al., 2023; Samaha and Postle, 2015; Shen et al., 2019; Wutz et al., 2018), the role of phase in the integration process remains unclear. Previous studies on the influence of oscillatory phase on perception primarily analysed absolute phase angles within a cycle, categorizing specific phases as favourable or unfavourable for perception (Busch et al., 2009; Mathewson et al., 2009; Neuling et al., 2012; Ng et al., 2012). This framework, however, creates a potential contradiction: one stimulus could be presented at a favourable phase and another at an unfavourable phase, leading to the erroneous perception of only one stimulus, irrespective of whether they occur in the same or different cycles. Such an explanation conflicts with our behavioural data, which suggest a more nuanced interplay between frequency and phase. Related work has also identified discrete perceptual cycles in the somatosensory domain (Baumgarten et al., 2015). Whether phase effects from near-threshold detection or another sensory modality generalize to the present suprathreshold double-flash judgments remains uncertain. Phase quality alone therefore cannot fully account for perception.

In contrast, our model assigns distinct roles to frequency and phase. Frequency determines whether the two flashes fall within the same alpha cycle, whereas phase modulates the reliability of the resulting report. This distinction may help interpret the null IAF effect reported by Buergers and Noppeney (2022), who assessed sensitivity and bias for two-flash discrimination rather than a fusion-threshold measure. However, it does not fully reconcile the divergent findings across audiovisual paradigms. Differences in sample size, stimulus and task design, psychometric measures, and IAF estimation may also contribute (Samaha and Romei, 2024). We therefore regard this model-based account as one possible explanation specific to the present two-flash fusion paradigm.

Our findings offer a novel perspective on phase resetting, emphasizing its influence on cross-modal interactions beyond its traditional role in facilitating multisensory integration. For example, cross-modal phase resetting can reorganize low-frequency activity in auditory cortex and has been linked to auditory task performance (Mégevand et al., 2020; Thorne et al., 2011). Similarly, auditory-driven phase resetting in the visual cortex has been demonstrated to facilitate audiovisual integration (Mercier et al., 2013). These studies highlight the critical role of phase resetting in optimizing sensory processing; its underlying mechanism involves the reorganization of neural oscillations through input from another sensory domain (Senkowski and Engel, 2024). Supporting this, a study in non-human primates by Lakatos et al. (2007) found that somatosensory inputs reset oscillatory phases in the auditory cortex by modulating subthreshold membrane potentials, rather than increasing neural firing rates, thereby enhancing responsiveness (Lakatos et al., 2007). The present findings suggest another possible consequence of such reorganization: auditory input may extend the temporal interval over which visual events are integrated. Although this wider window reduces temporal resolution for successive visual events, it may support cross-modal coherence when signals from different senses need to be combined. Phase resetting may therefore adapt the temporal properties of a sensory system to the current multisensory context.

The model was designed to test whether an auditory event could account for the F2B1−F2 difference when visual input was held constant. In the simulation, the phase trajectory was undelayed in F2 and delayed in F2B1; because visual stimulation was identical in the two conditions, no separate visual-reset parameter was included. The model therefore isolates the auditory contribution but does not establish the underlying physiological mechanism. The higher poststimulus ITC in F2B1 is consistent with phase resetting but could also reflect stimulus-evoked phase locking, which scalp EEG cannot distinguish. Intracranial recordings would provide a more direct test of this distinction (Bauer et al., 2020; Mégevand et al., 2020). The tACS experiment used fixed 8- and 13-Hz stimulation in an independent sample without individual alpha-frequency estimates, which may have limited sensitivity to small effects or effects that emerge only when stimulation is matched to each participant’s alpha frequency. tACS matched to individual alpha frequency would provide a stronger test of frequency-specific causality, while participant-level model fitting could test whether individual phase- delay or report-reliability parameters predict the F2B1−F2 shift in fusion threshold.

However, the present results do not identify where the auditory effect enters the proposed pathway. Stimulation over the occipital scalp site Oz targets the putative downstream visual alpha activity; a change in fusion threshold would test whether altering this activity is sufficient to affect temporal integration. Stimulation of auditory cortex would address the complementary upstream question of whether the beep-induced effect depends on auditory-cortex engagement. Together, these manipulations would distinguish the sufficiency of the downstream visual oscillator from the necessity of auditory-cortex engagement.

Our study shows that a concurrent sound can reshape how two successive visual events are integrated over time. Auditory input prolonged the behavioural fusion window, was associated with a transient slowing of poststimulus occipital alpha activity, and altered how alpha frequency and phase predicted perceptual outcomes. A phase-resetting model qualitatively reproduced these condition-dependent patterns, supporting a candidate account in which auditory input delays the visual alpha phase trajectory and transiently lengthens the ongoing cycle. Together, these findings suggest that the temporal window of visual integration is not fixed, but can be dynamically retuned by multisensory context through changes in alpha dynamics.

## Materials and methods

### Participants

Thirty-four healthy right-handed participants (23 females; age 20.8 ± 1.9, mean ± *SD*) participated in the EEG experiment. Another group of 30 participants participated in the tACS experiment (18 females; age 22.4 ± 2.2). All participants had normal or corrected-to-normal visual acuity and no history of neurological or psychiatric disease.

### Stimuli

The visual stimuli consisted of two identical white circles, each with a radius of 2 degrees, presented against a grey background. These circles were displayed rapidly one after the other, positioned peripherally below the fixation point at an eccentricity of 5 degrees of visual angle. Each visual stimulus had a duration of 10 milliseconds. The auditory stimulus was a 10- millisecond pure tone with a frequency of 3000 Hz and an approximate sound level of 50 decibels. To provide a spatially congruent experience for the participants, the auditory stimuli were delivered through a pair of loudspeakers situated on either side of the monitor. This arrangement created the perception that the auditory cues originated from the centre of the screen, closely aligned with the visual stimuli. Visual stimuli were presented using Presentation software (Neurobehavioural Systems; http://www.neurobs.com) on a 24-inch high-refresh-rate LCD monitor (resolution: 1920 × 1080, refresh rate: 100 Hz, viewing distance 57 cm) and viewed through a chin-and-forehead rest. The monitor was carefully calibrated using an oscilloscope before the experiment. The study consisted of two conditions: the F2 condition, involving the presentation of two visual stimuli, and the F2B1 condition, where the first visual flash was consistently accompanied by an auditory beep.

### Psychophysical pretest and threshold estimation

To determine each participant’s fusion threshold for the bistable condition, each participant completed a psychophysical pretest before the main experiment. Two physical flashes were presented in every trial, including both the 30-ms short-ISI and 100-ms long-ISI trials. During the formal psychophysical pretest, the first flash was presented for a duration of 10 ms; after a variable ISI (eight levels: 30, 40, 50, 60, 70, 80, 90, or 100 ms), the second flash was presented for 10 ms as well, either with no beep (F2) or with a beep accompanying the first flash (F2B1). Each ISI comprised 35 trials for both conditions. Participants were then asked to perform a two-alternative forced-choice task in which they had to choose between perceiving one flash or two flashes. Prior to the formal pretest, participants completed a practice block consisting solely of explicit short (30 ms) and long (100 ms) trials until accuracy reached at least 90%; a one-flash report was scored as correct for the 30-ms trials and a two-flash report for the 100- ms trials.

The same two-parameter logistic model was used for the pretest and tACS experiment. For a given ISI x, the probability of a two-flash report was defined as (Equation 1):

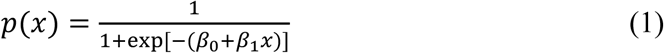

Here, β₀ is the intercept and β₁ is the change in log odds per millisecond. The binomial model was fitted by maximum likelihood with MATLAB glmfit. Setting the response probability to 0.5 gives the fusion threshold (Equation 2):

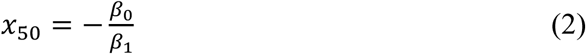

Because the outcome is a binomial count, the fitting criterion was the binomial likelihood rather than ordinary least-squares *R*^²^ (Wichmann and Hill, 2001). In the pretest, F2 and F2B1 were fitted separately using 35 trials per ISI.

### EEG experimental procedures

Participants maintained central fixation throughout the experiment. On each trial, they indicated whether they perceived one or two flashes by pressing one of two designated keys with the index or middle finger of their right hand. The mapping between the keys and the two perceptual reports was counterbalanced across participants. Because all critical comparisons were within-participant contrasts between the two sensory conditions, a response criterion that remained constant across conditions could not explain a difference in reports between conditions (Green and Swets, 1966; Knotts and Shams, 2016). However, a condition-specific shift in response criterion cannot be excluded. The task was organized into ten blocks with short breaks between blocks. The first nine participants were scheduled to complete 520 trials each, including 80 explicit short-ISI trials (30 ms; 40 F2 and 40 F2B1), 80 explicit long-ISI trials (100 ms; 40 F2 and 40 F2B1), and 360 bistable trials at the participant-specific fusion- threshold ISI (180 F2 and 180 F2B1). The remaining 25 participants were scheduled to complete 560 trials each, including 80 explicit short-ISI trials, 80 explicit long-ISI trials, and 400 bistable trials (200 F2 and 200 F2B1). Thus, F2 and F2B1 trials were equally represented within each planned trial type. Each trial began with 500–1,500 ms of central fixation, followed by a 10-ms presentation of the first visual stimulus. In the F2B1 condition, the first flash was accompanied by an auditory beep. After a variable ISI (30 ms, 100 ms, or the participant’s individual fusion threshold), the second flash was presented for 10 ms. A fixed 2,000-ms response period then followed, after which fixation for the next trial began.

### Recording and preprocessing of EEG data

The electroencephalogram (EEG) was recorded at a sampling rate of 1,000 Hz using a Neuroscan SynAmps2 system equipped with 64-channel Quick-Caps and Ag/AgCl electrodes. To capture horizontal and vertical electrooculograms, four extra electrodes were placed around the participants’ eyes. The impedances of all electrodes were kept below 5 kΩ. The signals were online-referenced to an electrode positioned between Cz and CPz. Continuous data were demeaned and linearly detrended; bad channels identified in individual participants were interpolated using a weighted average of neighbouring electrodes, and the signals were rereferenced offline to the average of the EEG electrodes. Data were filtered with a 49.5–50.5 Hz band-stop filter to eliminate 50-Hz line noise. No high-pass, low-pass, or broadband band- pass filter was applied during preprocessing. An 8–13 Hz least-squares finite-impulse-response filter, applied forward and reverse, was used only for the Hilbert-based alpha analyses. Data were epoched from −1,300 to 1,000 ms relative to first-flash onset and baseline-corrected using the mean amplitude over −500 to 0 ms. All inferential prestimulus analyses (instantaneous frequency, POS, phase-bin, and their baselines) were confined to the −600 ms immediately preceding stimulus onset. Signals containing blinks, muscular movements, and other EEG artifacts were detected and removed through independent component analysis. The data were then visually inspected, and trials containing residual artifacts or excessive noise were excluded. After artifact rejection, the data were downsampled to 500 Hz. Explicit short- and long-ISI analyses retained all artifact-free trials. The primary bistable IAF and POS-map analyses matched trial numbers across the four sensory-condition × report cells within each participant by randomly subsampling each cell to the smallest of the four cell counts.

### tACS experimental procedures

The alternating-current stimulation was administered using a 1 × 1 transcranial electrical stimulator (Soterix Medical, New York, NY) with rubber electrodes embedded in saline-soaked sponges. Electrodes were secured to the scalp with elastic bands. The reference and stimulation electrodes were positioned at the vertex (Cz) and posterior occipital area (Oz), respectively; montage and anatomical factors influence the resulting electric-field distribution (Opitz et al., 2015). Each electrode measured 5 × 7 cm². A sinusoidal current was used with a DC offset of 0, and impedance was maintained below 5 kΩ. The current intensity was set at 1.5 mA. Participants completed three 16-minute tACS sessions in randomised order: (1) 8 Hz stimulation (low-alpha), (2) 13 Hz stimulation (high-alpha), and (3) sham stimulation. Sessions were spaced 30 minutes apart. During sham stimulation, the current was ramped up over 30 seconds, then ramped down to zero over the subsequent 30 seconds, after which no further stimulation was applied. The Auto-Sham setting on the tACS controller was used to deliver sham stimulation. This ramping protocol mimicked the somatosensory sensation typically experienced during real transcranial electrical stimulation, particularly during current ramp-up or ramp-down phases. In each session, participants performed the same task as in the psychophysical pretest, with 20 trials for each ISI condition. Fusion thresholds were estimated separately for each stimulation session and sensory condition using the psychometric procedure described above.

### Alpha oscillation analysis of EEG data

The EEG analyses focused primarily on the bistable trials within both the F2 and F2B1 conditions, unless stated otherwise. We applied a fast Fourier transform (FFT) to obtain 1–30-Hz power spectra for all electrodes and participants. In each trial, the power spectrum was computed over the entire period of interest (−600 to 600 ms, relative to the presentation of the first flash) using a Hanning taper. The 1.2-s segments were zero-padded to 10 s, yielding a frequency-bin spacing of 0.1 Hz without increasing the intrinsic spectral resolution of the 1.2- s segment. Trial-level spectra were averaged across all bistable trials and all electrodes, and each participant’s alpha peak was defined as the frequency with maximal power between 8 and 13 Hz. Subsequent analyses used the 8–13 Hz band and the posterior region of interest defined below. The same maximum-power criterion was applied to 0–400 ms poststimulus spectra (F2 versus F2B1) and −400 to 400 ms peri-stimulus spectra for one-flash versus two-flash reports; these windows were chosen to match the significant intervals identified in the instantaneous-frequency analysis.

The Hilbert transform was then used to obtain the unwrapped alpha phase, *φ_u_*(*t*) . Instantaneous frequency was defined from its temporal derivative (Equation 3):

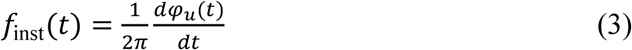

In Equation 3, phase is measured in radians; differentiation gives radians per second, and division by 2*π* converts the rate to hertz (Boashash, 1992; Cohen, 2014a). Estimates were retained every 10 ms. Because differentiation amplifies high-frequency noise, local medians were calculated within each trial in 10 centred windows with total widths from 10 to 400 ms, and the final value was the median across window-specific medians. Values were averaged over PO7, PO5, PO3, POz, PO4, PO6, PO8, O1, Oz, and O2. A lower value denotes slower local phase progression, not necessarily a sustained change in trait-like individual alpha frequency.

To account for the potential effects of the 1/f slope, we reconstructed the time-domain signals that had been demodulated by implementing 1/f attenuation, following the methods outlined in a previous study (Samaha and Cohen, 2022). Specifically, we used the Fast Fourier Transform (FFT) to extract the amplitude and phase spectra from each trial of the original signal. Second-order polynomials were fitted to the log10 amplitude spectra as a function of log10 frequency.

The log10 amplitude spectrum was divided point-by-point by the fitted values and then transformed back to the linear amplitude scale. The resulting amplitude spectrum was combined with the original phase spectrum, and an inverse FFT was used to reconstruct the time-domain signal. Next, the demodulated signal underwent a filter with a fixed frequency limit of 8–13 Hz to perform the instantaneous frequency analysis.

For the control analysis of instantaneous alpha power, we used the same filtering and Hilbert transform as in the IAF analysis to extract instantaneous alpha amplitude. The power was then calculated by squaring the amplitude.

Our primary goal was to investigate whether single-trial perception is influenced by the phase of ongoing oscillatory activity. To achieve this, we used the phase opposition sum (POS) method (VanRullen, 2016a) to evaluate the phase difference between the two distinct perceptual outcomes in the F2 and F2B1 conditions, using the common balanced bistable-trial set and the posterior region of interest comprising PO7, PO5, PO3, POz, PO4, PO6, PO8, O1, Oz, and O2. Because phase-concentration estimates depend on trial count, unequal 1-flash and 2-flash trial counts can make the two outcome estimates less comparable (Vinck et al., 2010). We therefore balanced the trial counts for both perceptual outcomes in both F2 and F2B1 conditions; thus, equal numbers of F2B1 one-flash, F2B1 two-flash, F2 one-flash, and F2 two- flash trials contributed within each participant.

Time-frequency transformations were first generated over occipital channels using FieldTrip (Oostenveld et al., 2011). Specifically, we used wavelets with logarithmically spaced cycles (ranging from 2 at the lowest frequency to 7 at the highest frequency) to obtain time-frequency representations (complex numbers) at 60 log-spaced frequency points spanning from 2 to 50 Hz, with time points from −600 to 400 ms in 20-ms steps and 10-s zero-padding. The reported POS maps were evaluated from −600 to 200 ms. A temporal-smearing control repeated the complete analysis with one-cycle wavelets at every frequency; for this control, the half-cycle boundary before stimulus onset, −1/(2f), was displayed on the time-frequency map to identify estimates whose wavelet support could extend beyond time zero (Figure 4—figure supplement 1).

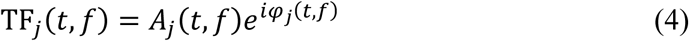

At each time-frequency point, the complex coefficient for trial *j* was written as in Equation 4. Dividing it by its magnitude retained phase while removing amplitude (Equation 5).

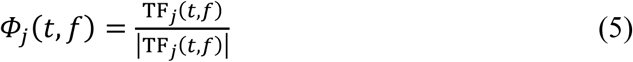

For either perceptual-report group containing *N* trials, inter-trial phase coherence was the length of the mean unit vector (Equation 6), as defined by Cohen (2014b). ITC ranges from 0 to 1 and measures phase concentration, not phase angle.

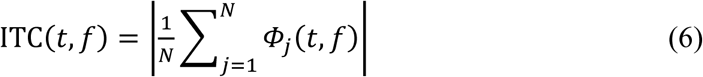

POS contrasted phase concentration in the one-flash and two-flash groups against the concentration obtained after pooling the groups (Equation 7), as defined by VanRullen (2016a).

Let *A* and *B* denote the one-flash and two-flash trial groups and let *all* denote their pooled trials:

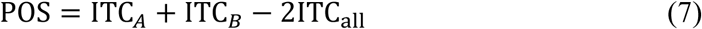

A large positive POS occurs when A and B are each phase concentrated but around different angles; if they cluster around the same angle, the pooled ITC also becomes large and POS approaches zero. This definition separates phase opposition from a nonspecific increase in phase locking.

For the phase-opposition control comparing high- and low-power trials, balanced bistable trials were split within each participant, sensory condition, and perceptual-report category by the median posterior prestimulus alpha power averaged from −600 to 0 ms. Phase opposition was recomputed separately for the high- and low-power subsets, and high-minus-low maps were evaluated with 100,000 within-participant label permutations. Cluster-based correction was restricted to frequencies ≥5 Hz to avoid low-frequency edge artifacts.

For the aligned phase-bin analysis, phase angles at 10.28 Hz, −300 ms, and Oz were divided into nine bins, and the proportion of two-flash reports was calculated for each bin. All artifact-free bistable trials were retained because this analysis estimates a within-participant phase- report function rather than a condition mean; retaining all trials stabilizes the bin-wise report proportions without changing the definition of the relative-phase profile. Each participant’s profile was circularly shifted so that the bin with the highest two-flash proportion occupied the centre. The central bin was excluded from inferential testing because its maximum was imposed by this alignment. Quadratic contrasts across the remaining eight relative-phase bins tested the phase-bin effect and its interaction with sensory condition. These aligned response probabilities are continuous dependent variables indexed by relative phase distance; they are not a sample of raw angular observations to which a Watson-Williams test can be directly applied. The participant-level mean phase directions were instead compared with Watson– Williams tests (Berens, 2009).

As a supplementary non-aligned measure of phase-dependent modulation, we extracted phase angles at the illustrative point (10.28 Hz, −300 ms, Oz) from all artifact-free bistable trials, retaining all valid trials because this analysis estimates each participant’s within-condition phase-response profile rather than a condition mean. Phase angles were divided into *K* = 12 equally spaced bins, and the proportion of two-flash reports in each bin was normalized to sum to one, yielding a phase-response profile. We quantified the strength of phase-dependent modulation as the Kullback–Leibler (KL) divergence of this profile from a uniform profile (Equation 8):

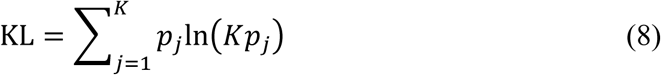

Here, *p_j_* is the normalized two-flash-report probability in phase bin *j*. A uniform profile assigns probability 1/K to every bin, so *Kp_j_* is the observed-to-uniform ratio: KL is zero for a uniform profile and increases as two-flash reports concentrate near a preferred phase. To test whether the observed KL exceeded chance, response labels were shuffled 10,000 times within each participant and condition, preserving the number of trials in each report category, and KL was recomputed for each shuffle to form a participant-specific null distribution. The observed KL was standardized against this null distribution as *z* = (KL − *μ*)/*σ*, where *μ* and *σ* are the mean and standard deviation of the shuffled KL values. Within each condition, the standardized scores were tested against zero with right-tailed one-sample and sign-flip tests; F2B1 and F2 were compared with a two-tailed paired test and a sign-flip test (Kullback and Leibler, 1951; Tort et al., 2010).

### Statistical analyses

Behavioural comparisons used two-tailed paired-samples *t* tests unless a directional contrast had been specified a priori. For the tACS experiment, directional contrasts were specified within each sensory condition: because slower alpha was predicted to widen the temporal integration window, 8-Hz stimulation was expected to yield a higher fusion threshold than 13- Hz stimulation. These contrasts were evaluated one-tailed and corrected across the two conditions with the Benjamini–Hochberg false-discovery-rate (FDR) procedure; sham comparisons, for which no direction was predicted, were evaluated two-tailed and reported descriptively. Whenever a family of follow-up tests addressed the same hypothesis, *p* values were FDR-corrected at *q* = .05, and both uncorrected and corrected values are reported. For the KL summary analysis, the two condition-specific tests and their paired comparison formed one FDR family. Before parametric tests, the relevant within-participant difference scores were screened with Shapiro–Wilk tests and Q–Q plots; when normality was rejected, the parametric result is accompanied by a Wilcoxon signed-rank test and a bias-corrected bootstrap confidence interval on the mean difference (10,000 resamples), and a conclusion is drawn only where these agree. This screening was not applied to permutation, circular, or cluster-based tests, whose inference does not rest on a parametric normality assumption. Correlations used Pearson’s *r*, and Bayes factors used a default Cauchy prior (*r* = 0.707), two-sided for all reported Bayes factors, including the tACS contrasts. Time-resolved EEG analyses used cluster-based permutation correction (Maris and Oostenveld, 2007). For poststimulus-versus-baseline tests, FDR correction was applied jointly across sensory conditions and trial types.

POS significance was assessed separately in F2B1 and F2 using a nonparametric label-permutation procedure. Within each participant, one-flash and two-flash labels were randomly reassigned while preserving the original group sizes. POS was then recomputed, averaged across posterior electrodes, and averaged across participants. One hundred thousand permutations formed the null distribution at each time-frequency point, and pointwise *p* values were calculated with a plus-one correction. Cluster correction for the condition-specific POS maps was applied over the prestimulus interval from −600 to 0 ms. The direct F2B1−F2 comparison summed POS over 8–13 Hz and −600 to 0 ms and compared the observed condition difference with the corresponding distribution of permutation differences. In the figures, dashed lines mark the percentile boundaries of the permutation null distributions (i.e., the boundaries of the two-tailed 5% critical region). The same sign-flip framework was used for the non-aligned KL analysis: participant-level KL *z* scores were tested against zero with right-tailed permutations within each condition, and the F2B1−F2 difference was tested with a two-tailed paired permutation test.

### Simulations of EEG data

The computational model was implemented in MATLAB (workflow in Figure 8—figure supplement 1). Simulated signals extended from −1.3 to 1.0 s at 1,000 Hz, with a 1-ms time step and 1,000 trials per condition. On each trial, oscillator frequency was sampled from a normal distribution with mean 10 Hz and *SD* 1 Hz, amplitude was sampled uniformly from 5 to 20, and initial phase was sampled uniformly from −π/2 to π/2. The original oscillator term was retained as (Equation 9):

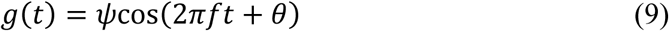

For each trial, the simulated signal was advanced using the original update equation (Equation 10):

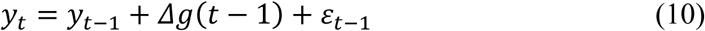

Here, the time step was 1 ms and the innovation term was independent Gaussian noise with *SD* 0.01, chosen because additive, uncorrelated process noise with no systematic drift is the simplest neutral perturbation. F2 used the undelayed update throughout each trial. In F2B1, perturbation onset was sampled uniformly from 0 to 50 ms after the first flash; from that sample onward, the oscillator term was evaluated with a fixed 25-ms phase delay, chosen because it lies within the 10–30 ms range across which the sensitivity analysis (Figure 8—figure supplements 3 and 4) reproduces the qualitative pattern, and because it yields a transient slowing on the order of the observed 0–280 ms window. Thus, onset jitter determines when the auditory perturbation begins, whereas the fixed delay determines its magnitude in time. The resulting signals were filtered from 8 to 13 Hz with a zero-phase fourth-order Butterworth filter, and analytic phase was extracted with the Hilbert transform.

Frequency and phase-delay sensitivity analyses repeated the simulation while holding the remaining pipeline fixed. The frequency analysis used fixed 8, 9, 10, 11, 12, and 13 Hz oscillations with a 25-ms delay; the delay analysis used delays of 10, 20, 30, and 40 ms with trial frequencies sampled from *N*(10, 1 Hz). Each parameter setting contained 1,000 simulated trials.

For each condition, 1,000 trials were evaluated at eight ISIs from 30 to 100 ms in 10-ms steps, including the 30-ms and 100-ms ISIs used for the explicit short- and long-ISI trials, respectively; the full set generated the simulated psychometric curves shown in Figure 8A. Alpha phase was sampled at the two flash times. Whether the flashes occupied the same or different cycle defined the latent one-flash/two-flash outcome. The phase of the second flash controlled report reliability: when the second flash fell within the favourable phase range (−*π*/2 to *π*/2), the report followed the latent cycle-based outcome; outside that range, the report was sampled with equal probability from the two alternatives to represent reduced perceptual clarity. Frequency and phase therefore play conceptually distinct roles: frequency determines the perceptual outcome (whether the two flashes are integrated into one percept or seen as separate), whereas phase determines perceptual clarity (how reliably that outcome is reported). A two-parameter logistic function was fitted to the simulated proportion of two-flash reports, and its 50% point defined the simulated fusion threshold. Subsequent frequency and phase analyses were performed at the condition-specific threshold. The model did not separately parameterize a shared visual phase effect for F2 and F2B1; possible audiovisual interactions are outside the current model’s scope.

To quantify how well simulated reports discriminated between the model-defined latent states, we used the model-based discriminability index *D_model_*, defined in Equation 11. The latent two-flash state was treated as signal present, and the latent one-flash state as signal absent.

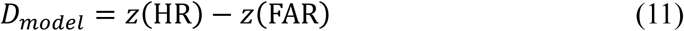

The hit rate (HR) was the proportion of latent two-flash trials reported as two flashes. The false-alarm rate (FAR) was the proportion of latent one-flash trials reported as two flashes. In Equation 11, *z* denotes the quantile (inverse cumulative distribution) function of the standard normal distribution; it converts HR and FAR into their corresponding standard-normal values. Thus, the resulting *D_model_* is a model-based index of how well simulated reports discriminate the model-defined latent states, not a conventional estimate of participants’ perceptual sensitivity.

Before the simulated EEG analyses, trials were randomly subsampled to the smallest of the four condition-by-report cells. Simulated instantaneous frequency and phase opposition were estimated with the same pipelines and time-frequency parameters as the empirical analyses (100,000 response-label permutations for the simulated POS).

## Additional information

### Author contributions

Mengting Xu: Conceptualization, Data curation, Formal analysis, Methodology, Visualisation, Project administration, Writing – original draft, Writing – review & editing. Biao Han: Conceptualization, Data curation, Formal analysis, Methodology, Visualisation, Supervision, Funding acquisition, Writing – review & editing. Qi Chen: Conceptualization, Supervision, Funding acquisition. Lu Shen: Conceptualization, Data curation, Formal analysis, Methodology, Visualisation, Supervision, Project administration, Funding acquisition, Writing – original draft, Writing – review & editing.

### Data and code availability

The data and analysis code are publicly available on the Open Science Framework: https://doi.org/10.17605/OSF.IO/2HSQW.

### Ethics

Human participants: The study was approved by the Ethics Committee of the School of Psychology, South China Normal University, and was conducted in accordance with the Declaration of Helsinki. All participants provided informed consent before participation.

## Supplementary Information

### Supplementary Results

#### Explicit-trial accuracy

At the two fixed ISIs, the auditory beep shifted report accuracy toward fusion. At the 30-ms ISI, where the expected report was one flash, the beep increased one-flash accuracy; at the 100-ms ISI, where the expected report was two flashes, the beep reduced two-flash accuracy. This produced a reliable sensory-condition × ISI interaction, *F*_(1, 33)_ = 12.08, *p* = .001 (Figure 1—figure supplement 3A). We next asked whether this behavioural shift tracked the auditory-induced change in oscillatory timing. The auditory-induced change in report accuracy (ΔAccuracy, F2B1 − F2) did not correlate with the auditory-induced change in poststimulus alpha-cycle length (ΔCycle length, F2B1 − F2) in the short-ISI trials (*r* = −.01, *p* = .946) or the long-ISI trials (*r* = −.08, *p* = .671, *n* = 34; Figure 1—figure supplement 3B and C). Thus, no significant association between these two auditory-induced changes was detected at either fixed ISI.

#### Instantaneous alpha frequency after 1/f removal

To test whether the F2 frequency–percept difference could be explained by the aperiodic (1/f) component of the spectrum rather than by the alpha rhythm itself, we repeated the analysis after attenuating the aperiodic component of every single-trial spectrum. The F2 frequency–percept difference remained similar in direction and magnitude (0.079 Hz, *t*_(33)_ = 1.86, one-tailed uncorrected *p* = .036, FDR-corrected *p* = .072), but did not survive correction; the condition × percept interaction was also not significant, *F*_(1, 33)_ = 0.64, *p* = .430 (Figure 3—figure supplement 1C). The anti-1/f procedure compresses the alpha peak relative to neighbouring frequencies, and the standard error was correspondingly larger than in the primary analysis (0.043 vs. 0.023 Hz). The F2 frequency–percept difference therefore remained similar in direction and magnitude after 1/f removal, consistent with the pattern observed in the primary analysis.

#### Poststimulus frequency decreases in explicit trials

In the explicit short- and long-ISI trials, F2 and F2B1 shared identical visual sequences. Poststimulus instantaneous alpha frequency was lower in F2B1 than in F2 over 0–180 ms at the short ISI (*p* = .035) and 0–370 ms at the long ISI (*p* = .001; Figure 2—figure supplement 1A and B). We also compared mean frequency over 0–600 ms with the −600 to −10 ms baseline. F2B1 showed reliable decreases at both the short ISI (Δ = −0.051 Hz, *t*_(33)_ = −2.01, *p* = .026, FDR-corrected *p* = .039) and the long ISI (Δ = −0.115 Hz, *t*_(33)_ = −3.62, *p* < .001, FDR-corrected *p* = .003). In F2, the short-ISI decrease did not reach significance (Δ = −0.039 Hz, *t*_(33)_ = −1.41, *p* = .084, FDR-corrected *p* = .101), whereas the long-ISI decrease was significant (Δ = −0.076 Hz, *t*_(33)_ = −2.82, *p* = .004, FDR-corrected *p* = .012; Figure 2—figure supplement 1C). All baseline tests were one-tailed and FDR-corrected, following the predicted direction of a poststimulus slowing. The long-ISI F2 decrease indicates that the visual sequence itself can produce some poststimulus slowing, so the auditory contribution is best interpreted from the direct F2B1−F2 contrast.

#### Parameter sensitivity of the simulation

To test whether the main simulation’s conclusions depended on the chosen oscillator frequency and phase delay, we repeated the simulation while holding the remaining pipeline fixed. The frequency analysis used fixed 8, 9, 10, 11, 12, and 13 Hz oscillations with a 25-ms delay, and the delay analysis used delays of 10, 20, 30, and 40 ms with trial frequencies sampled from *N*(10, 1 Hz). Each parameter setting contained 1,000 simulated trials and was processed with the same analysis pipeline as the main simulation.

##### Sensitivity to oscillator frequency

For oscillator frequencies of 9–12 Hz, the simulated psychometric curves were monotonic with well-defined fusion thresholds, and the auditory perturbation produced a consistent rightward threshold shift in F2B1 relative to F2 (9 Hz: 54.6 vs. 76.1 ms; 10 Hz: 48.6 vs. 69.8 ms; 11 Hz: 44.7 vs. 61.7 ms; 12 Hz: 38.6 vs. 54.9 ms). At 8 Hz, the F2B1 curve became non-monotonic and its logistic threshold was undefined: because one cycle at 8 Hz (125 ms) is longer than the tested ISI range (30–100 ms), the phase reset displaced the phase of the second flash such that the cycle-based perceptual outcome reversed across ISIs. At 13 Hz, the F2 curve rolled over at long ISIs: one cycle at 13 Hz (77 ms) falls within the ISI range, so the two flashes could cross more than one full cycle and the two-flash report proportion declined beyond roughly 70 ms. The clean operating range therefore brackets 10 Hz, and the main simulation’s random frequency N(10, 1 Hz) keeps most trials within this range (Figure 8—figure supplement 3).

##### Sensitivity to phase delay

The fusion-threshold shift increased monotonically with the imposed delay (10 ms: 4.4 ms; 20 ms: 12.8 ms; 30 ms: 23.5 ms; 40 ms: 30.1 ms), consistent with the model’s mechanism, whereby delaying the phase trajectory extends the fusion window. The simulated poststimulus IAF decrease, however, was non-monotonic: it was robust for delays of 10–30 ms (dips of 0.39–0.70 Hz centred near stimulus onset) but essentially absent at 40 ms (dip of 0.04 Hz, with a slight poststimulus frequency increase). At 40 ms the imposed delay approaches half of the ∼10-Hz cycle; for a substantial fraction of trials the delayed phase reference advances past the oscillator’s current phase rather than lagging it, so phase-advance trials cancel the group-level frequency decrease. The main simulation’s 25-ms delay (∼0.25 cycle) lies within the robust regime (Figure 8—figure supplement 4).

These analyses support the parameter choice of the main simulation (trial frequencies sampled from N(10, 1 Hz) with a 25-ms auditory phase delay): within this regime the model produces monotonic psychometric functions, a consistent F2B1 threshold shift, and a transient poststimulus IAF decrease, as observed empirically.

## Figure supplements

**Figure 1—figure supplement 1.**
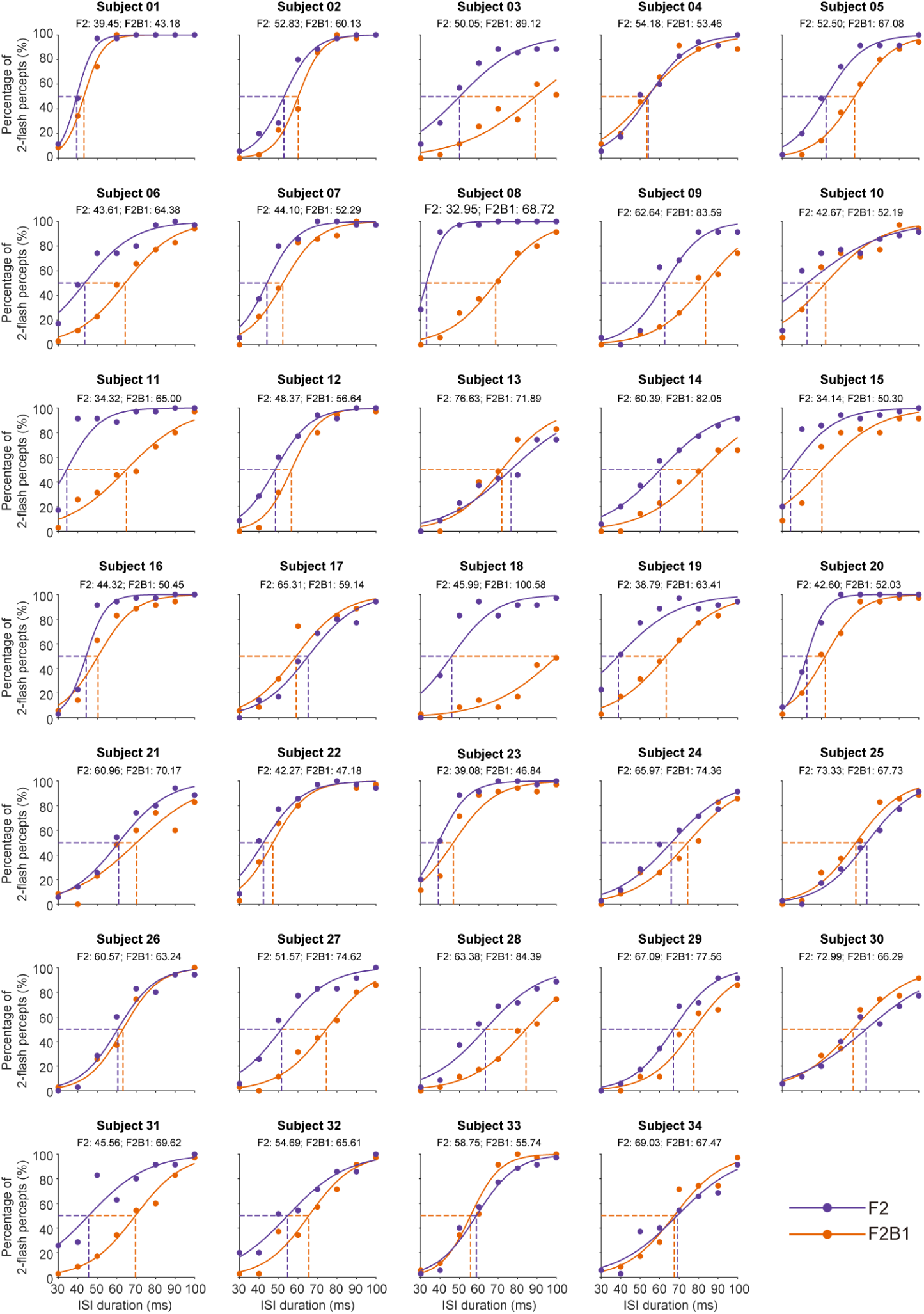
Behavioural data from the psychophysical pretest for each individual participant. Each panel shows one participant’s percentage of two-flash reports as a function of ISI for F2 (purple) and F2B1 (orange); symbols are the observed proportions, continuous lines the fitted binomial logistic curves, and vertical dashed lines the fitted 50% fusion thresholds, whose values are printed in each panel.

**Figure 1—figure supplement 2.**
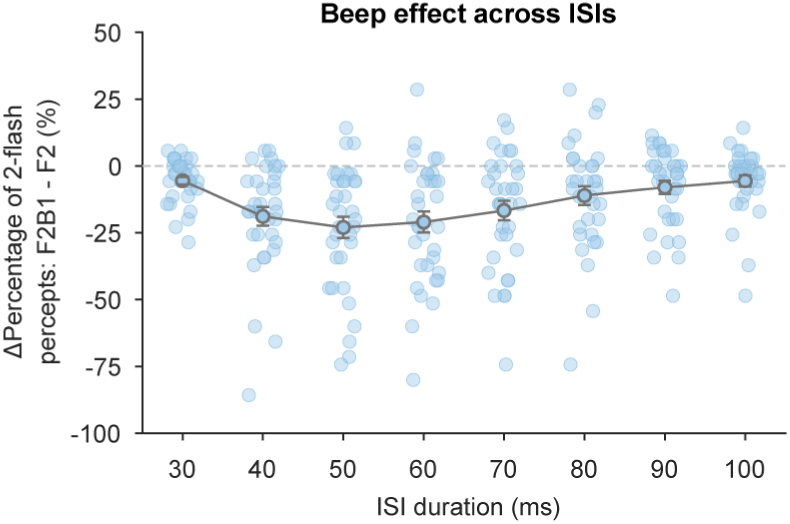
Differences in the proportion of two-flash reports between F2 and F2B1 across ISIs. Light blue dots show individual participants and the grey line the group mean ± *SEM* of the percentage difference (F2B1 − F2) at each ISI; the dashed line marks zero.

**Figure 1—figure supplement 3.**
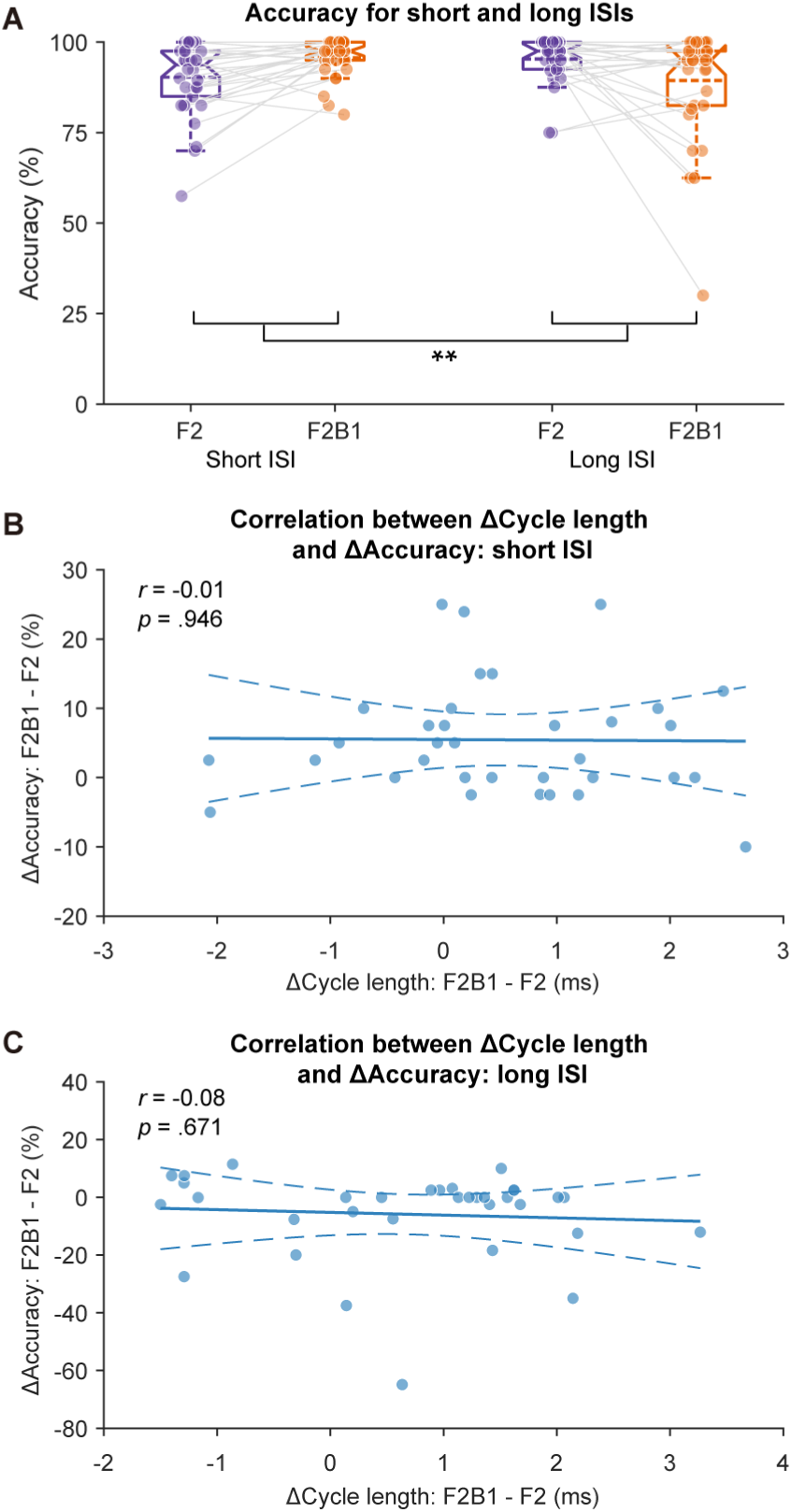
Explicit-trial behavioural results and their relation to alpha-cycle length. (A) Report accuracy at the two fixed ISIs: at the 30-ms ISI the expected report was one flash, and at the 100- ms ISI the expected report was two flashes; accuracy is plotted separately for F2 (purple) and F2B1 (orange), with dots for individual participants, boxes for the median and interquartile range, whiskers for the non-outlier range, dashed lines for the mean, and grey lines connecting the within-participant F2–F2B1 observations (ISI × sensory condition interaction: repeated-measures ANOVA). (B, C) Auditory-induced change in report accuracy (ΔAccuracy, F2B1 − F2) plotted against the auditory-induced change in poststimulus alpha-cycle length (ΔCycle length, F2B1 − F2), separately for the short-ISI (B) and long-ISI (C) trials; dots are individual participants and the solid line is the least-squares fit and dashed lines show its 95% confidence band, with the Pearson *r* and *p* values printed in each panel (uncorrected correlation *p* values). \*\**p* < .01.

**Figure 1—figure supplement 4.**
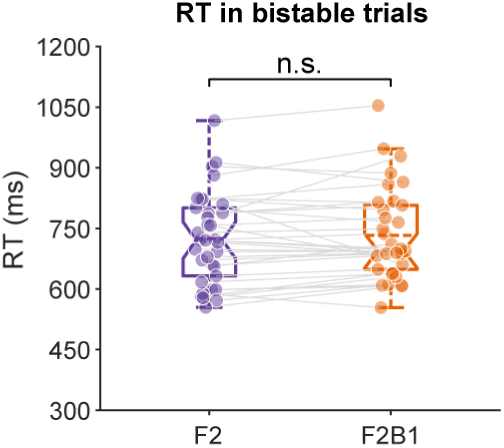
Reaction times in the bistable condition. Participant-level mean reaction times for F2 and F2B1 in the bistable trials, at which the perceptual outcome is maximally ambiguous. Boxes show the median and interquartile range, whiskers the non-outlier range, dots individual participants, and grey lines the within-participant pairing (uncorrected paired test). Reaction times at the fixed short and long ISIs, and the comparison between one- and two-flash reports, are not reported because those conditions were not designed to equate perceptual ambiguity. n.s., not significant.

**Figure 2—figure supplement 1.**
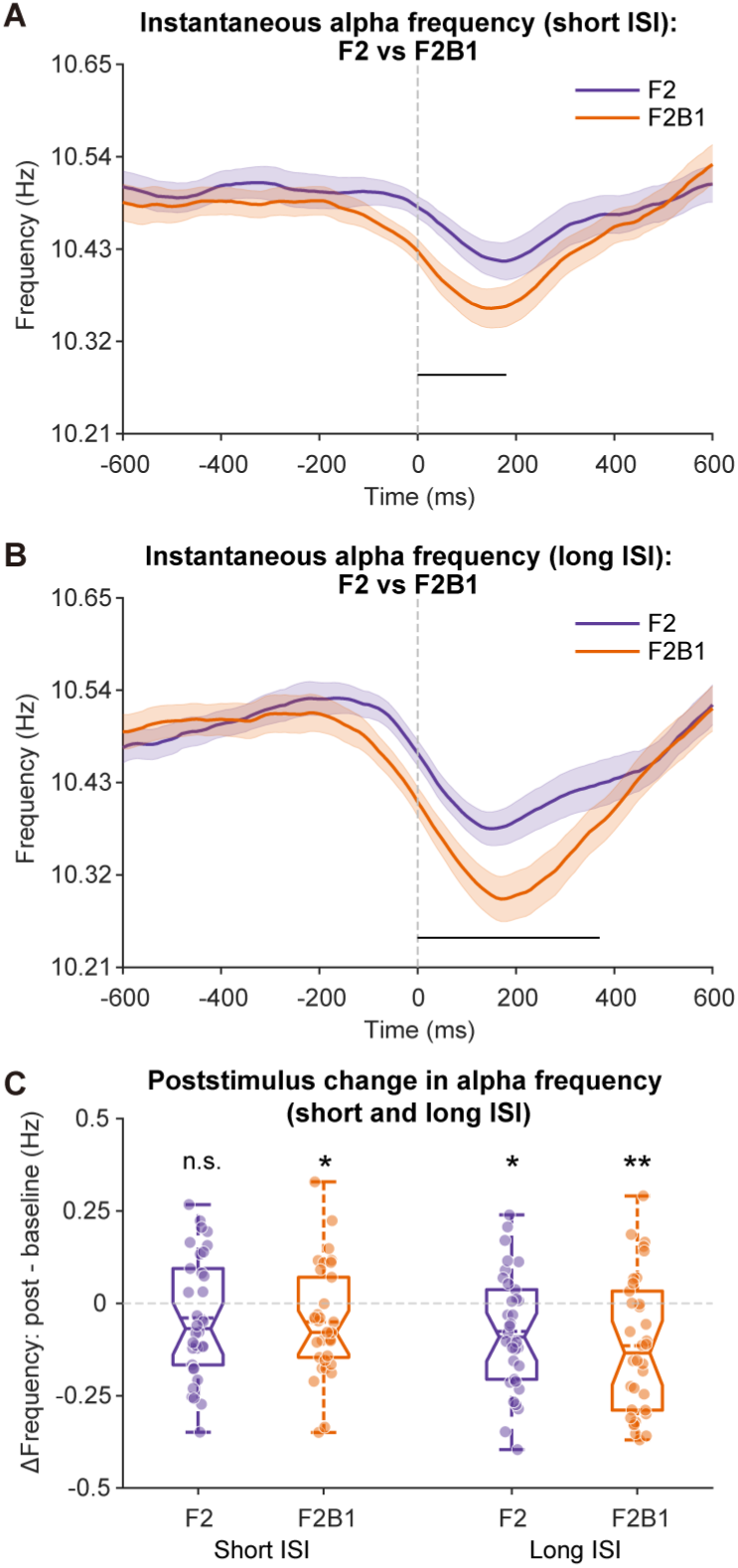
Instantaneous alpha frequency at the fixed physical ISIs. (A) Instantaneous alpha frequency for F2 (purple) and F2B1 (orange) in the short-ISI (30 ms) trials, averaged over the posterior region of interest; shaded areas show ±1 within-subjects *SEM* and the black horizontal bar marks the significant cluster (0–180 ms). (B) Similar to (A), but for the long-ISI (100 ms) trials (0–370 ms). (C) Instantaneous alpha frequency averaged over 0 to 600 ms minus the −600 to −10 ms baseline, for each fixed ISI and sensory condition. Boxes show the median and interquartile range, whiskers the non-outlier range, dashed lines the mean, and dots individual participants (FDR-adjusted one-tailed tests). \*\**p* < .01; \**p* < .05; n.s., not significant.

**Figure 2—figure supplement 2.**
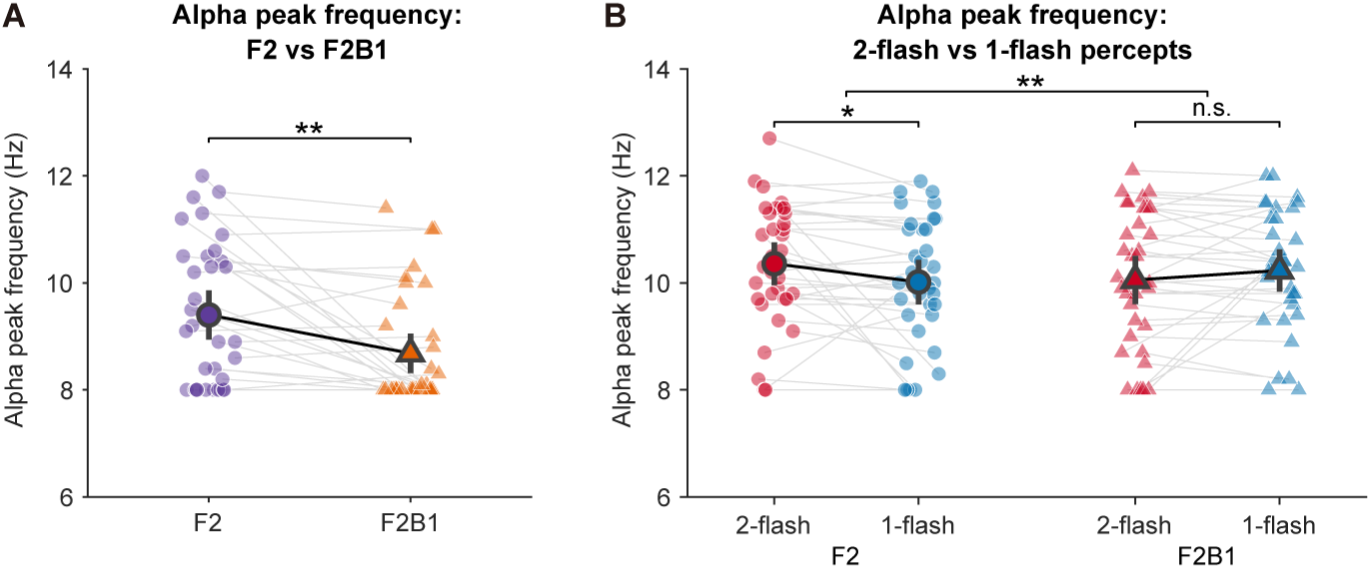
Alpha peak frequency estimated from the power spectrum. Peak frequency was taken as the 8–13 Hz bin with maximal power in the trial-averaged spectrum of the balanced bistable trials. (A) Poststimulus window (0 to 400 ms), F2 versus F2B1 (uncorrected two-tailed paired test). (B) Peri-stimulus window (−400 to 400 ms), two-flash versus one-flash reports in each condition (interaction: repeated-measures ANOVA; within-condition comparisons: uncorrected one-tailed paired tests). Error bars show 95% confidence intervals; dots show individual participants, and grey lines connect the paired observations within each comparison. Peak frequencies were read from a fixed 8–13 Hz search range. \*\**p* < .01; \**p* < .05; n.s., not significant.

**Figure 2—figure supplement 3.**
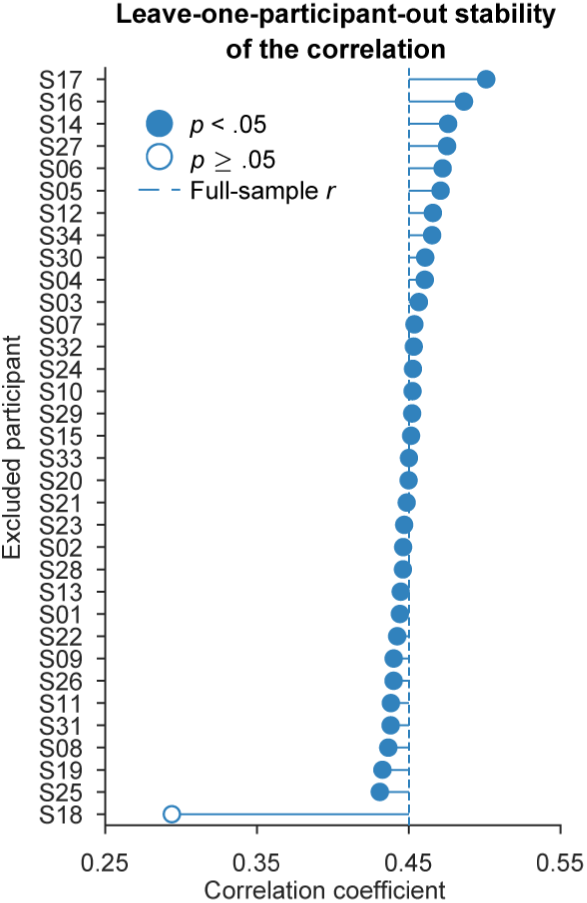
Robust association between the auditory-induced change in fusion threshold and poststimulus alpha-cycle length. Leave-one-participant-out Pearson correlations between the auditory-induced change in poststimulus alpha-cycle length (ΔCycle length, F2B1 − F2) and the auditory-induced change in the fusion threshold (Δfusion threshold, F2B1 − F2). Each row omits one participant; the dashed line marks the full-sample estimate. Uncorrected *p* values: filled markers, *p* < .05; open marker, *p* ≥ .05.

**Figure 3—figure supplement 1.**
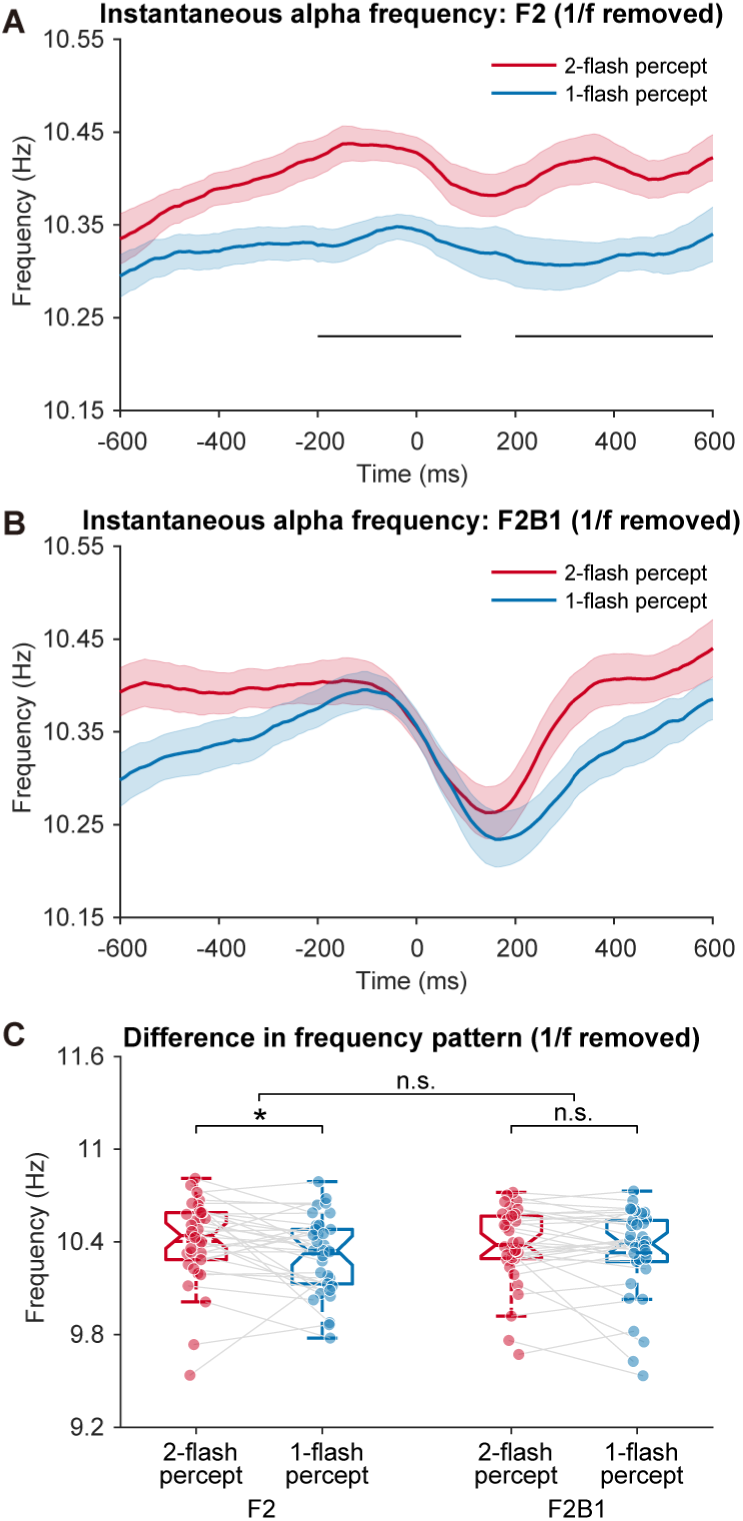
Instantaneous alpha frequency (IAF) after removal of the 1/f component. The instantaneous alpha frequency (IAF) analysis was repeated after attenuating the aperiodic component of every single-trial spectrum. (A) IAFs over time for two-flash (red) and one-flash (blue) reports in the F2 condition; black horizontal bars mark significant clusters. Shaded areas show ±1 within-subjects *SEM*. (B) Similar to (A), but for the F2B1 condition. (C) IAFs averaged over −600 to 600 ms for each condition and perceptual report. Boxes show the median and interquartile range, whiskers the non-outlier range, dashed lines the mean, and dots individual participants (interaction: uncorrected repeated-measures ANOVA; within-condition comparisons: uncorrected one-tailed paired tests). \**p* < .05; n.s., not significant.

**Figure 3—figure supplement 2.**
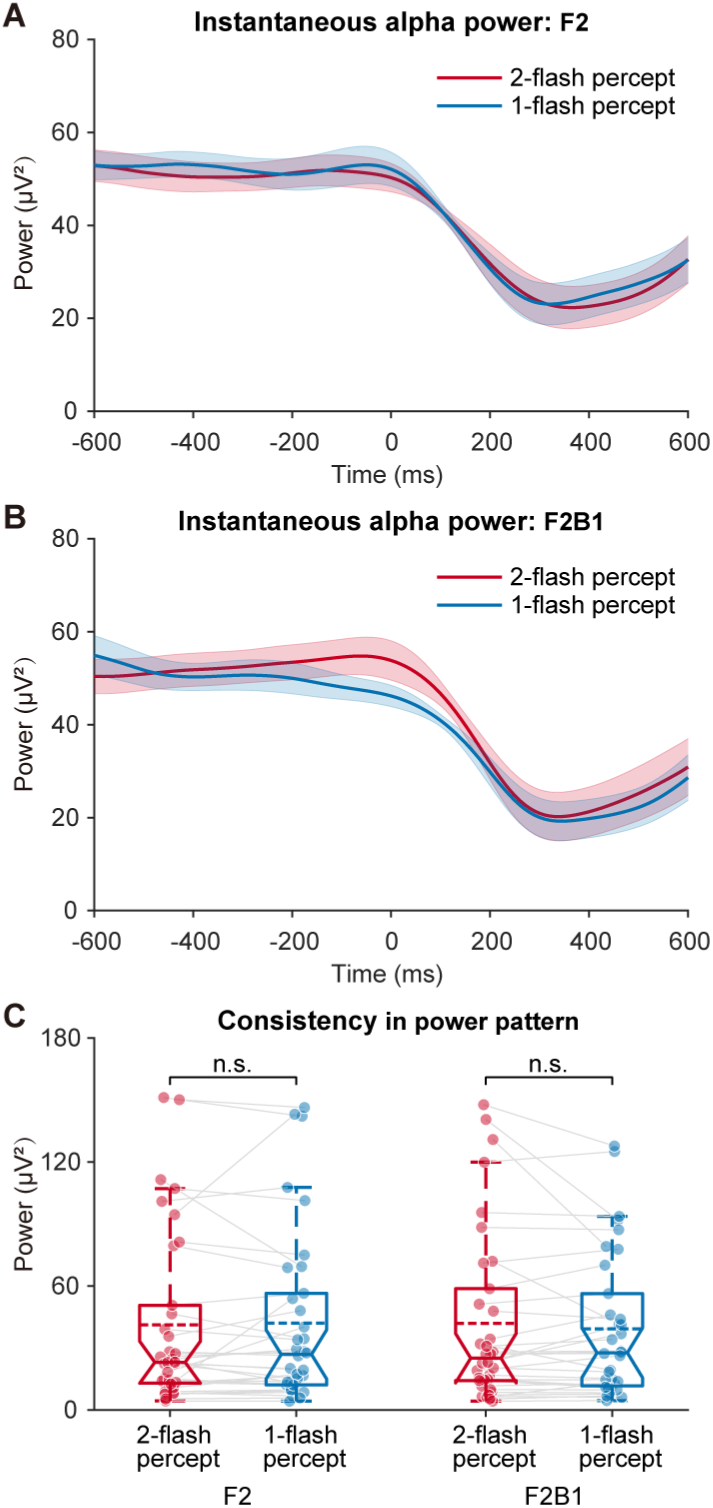
Relationship between the instantaneous alpha power (IAP) and perceptual outcomes. (A) IAPs over time for two-flash (red) and one-flash (blue) reports in the F2 condition; shaded areas show ±1 within-subjects *SEM*. (B) Similar to (A), but for the F2B1 condition. (C) IAPs during the time of interest (−600 to 600 ms) averaged separately for F2 and F2B1 conditions with different perceptual outcomes. Boxes show the median and interquartile range, whiskers the non-outlier range, dashed lines the mean, and dots individual participants (FDR-adjusted two-tailed paired tests). n.s., not significant.

**Figure 4—figure supplement 1.**
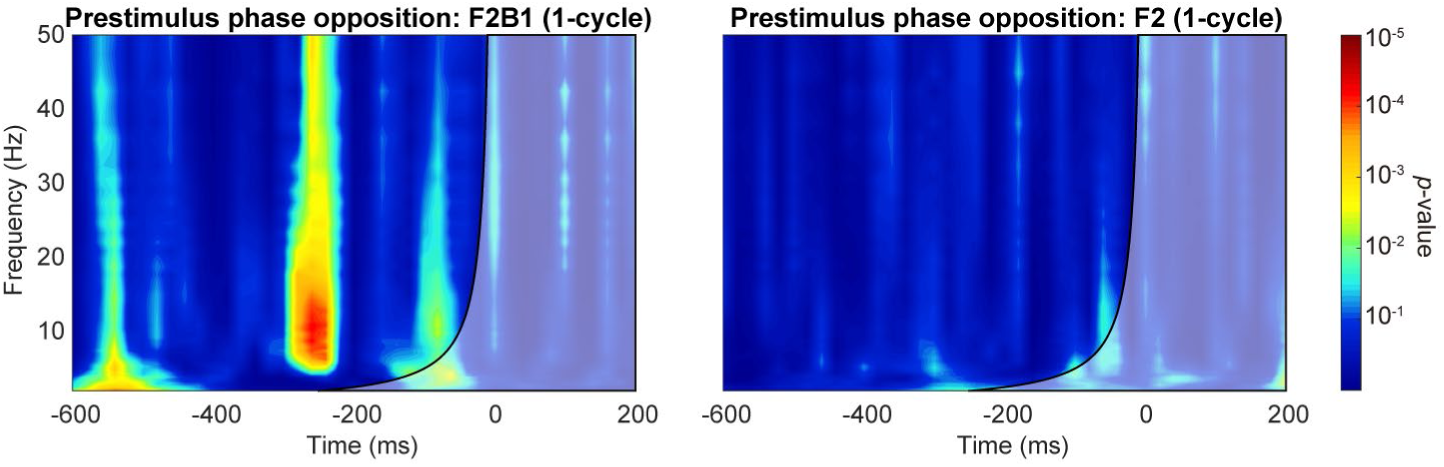
Temporal-smearing control for the prestimulus phase-opposition analysis. The prestimulus phase-opposition analysis was repeated with one-cycle wavelets at every frequency. Maps are shown for F2B1 (left) and F2 (right); colour encodes pointwise permutation −log10(*p*) (colour bar on the right). The black curve marks the frequency-specific half-cycle boundary −1/(2f), and the grey shading identifies estimates whose wavelet support could reach stimulus onset. The one-cycle control descriptively reproduced a prestimulus F2B1 alpha-band pattern without a comparable F2 pattern; no cluster-level inference was applied to this control.

**Figure 4—figure supplement 2.**
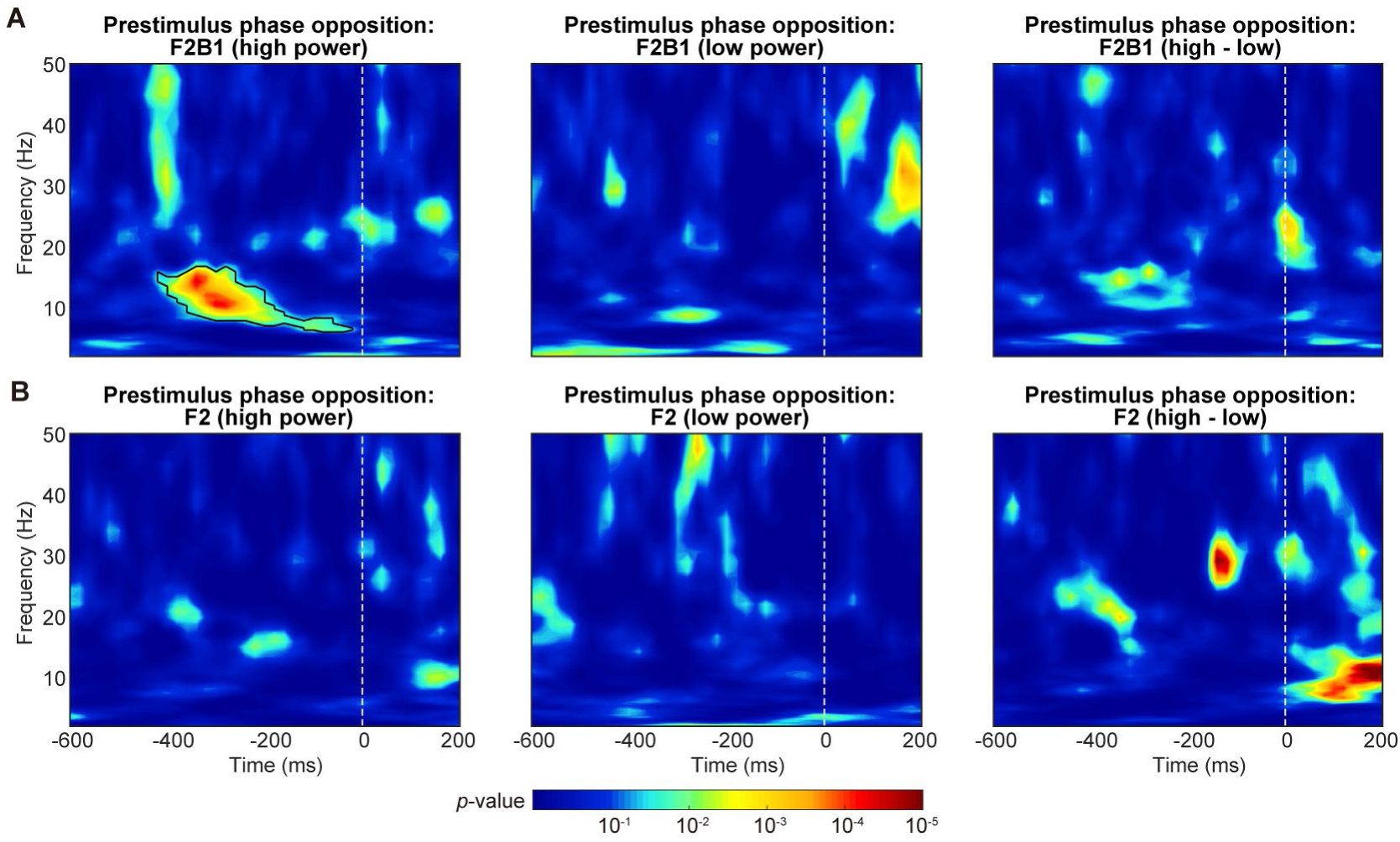
Phase opposition in high- versus low-power trials. POS was recomputed in high- and low-power subsets within each condition (median split by posterior prestimulus alpha power). (A) F2B1: high-power, low-power, and high-minus-low maps. (B) F2: high-power, low-power, and high-minus-low maps. Colour encodes −log10(*p*) (colour bar at the bottom) and black contours indicate clusters surviving permutation-based correction restricted to frequencies ≥5 Hz. This descriptive control does not itself test a formal condition × power interaction.

**Figure 4—figure supplement 3.**
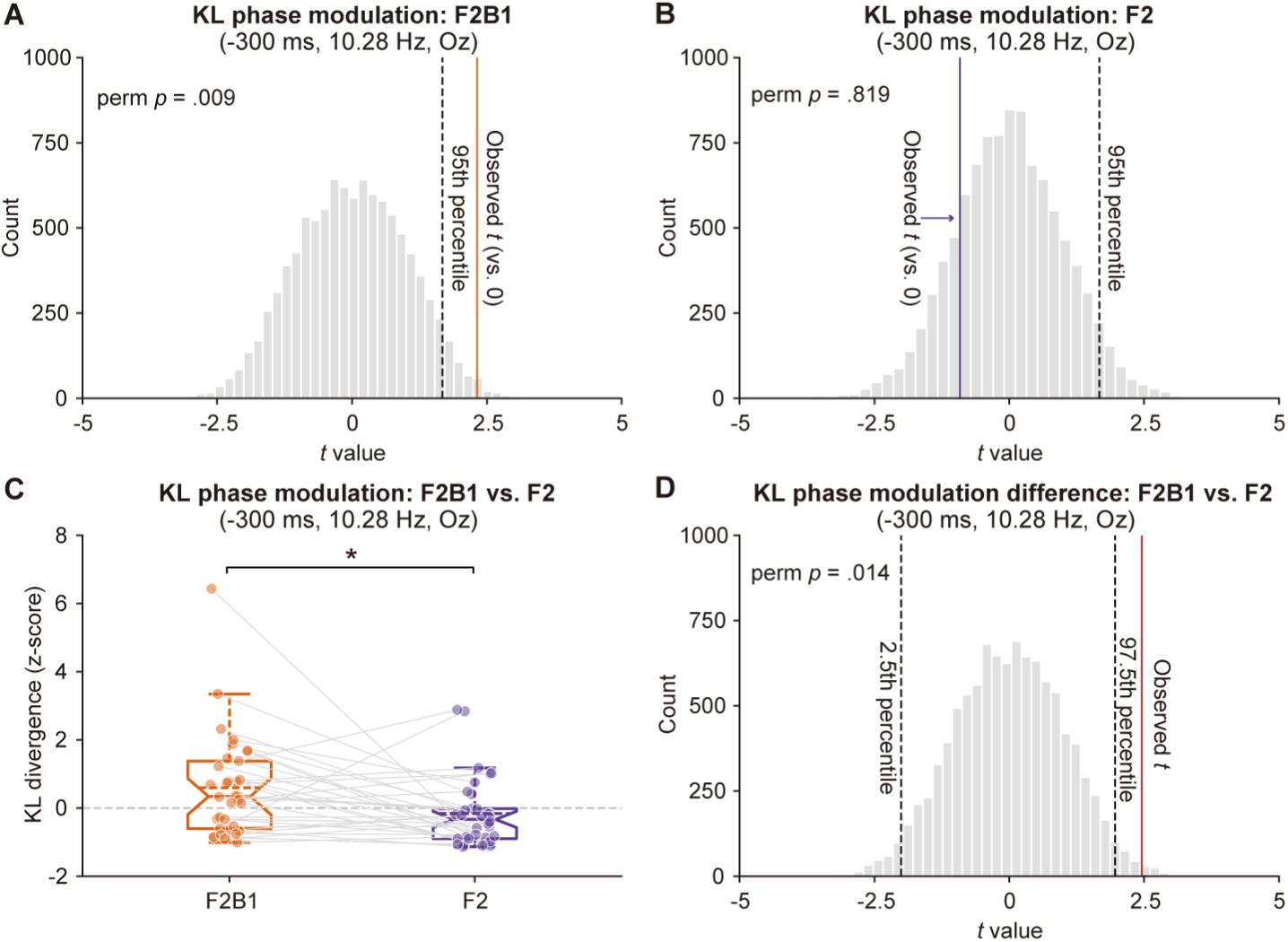
Non-aligned KL phase-modulation analysis. The non-aligned Kullback– Leibler (KL) divergence analysis quantifies each participant’s phase-dependent modulation of two-flash reports against a participant-specific response-label permutation distribution. (A) F2B1: the grey histogram shows the permutation distribution of *t* values, the orange line the observed *t* value, and the dashed black line the 95th percentile of the null distribution; the permutation *p* value is printed in the panel. (B) The same analysis for F2, with the observed line in purple. (C) Participant-level KL *z*-scores for F2B1 (orange) and F2 (purple): boxes show the median and interquartile range, whiskers the non-outlier range, dots individual participants, grey lines the within-participant pairing, and the dashed horizontal line marks zero (FDR-adjusted paired test). (D) Null distribution for the F2B1 − F2 difference: the grey histogram shows the permutation distribution of *t* values, the red line the observed *t* value, the dashed black lines the 2.5th and 97.5th percentiles, and the permutation *p* value is printed in the panel. \**p* < .05.

**Figure 5—figure supplement 1.**
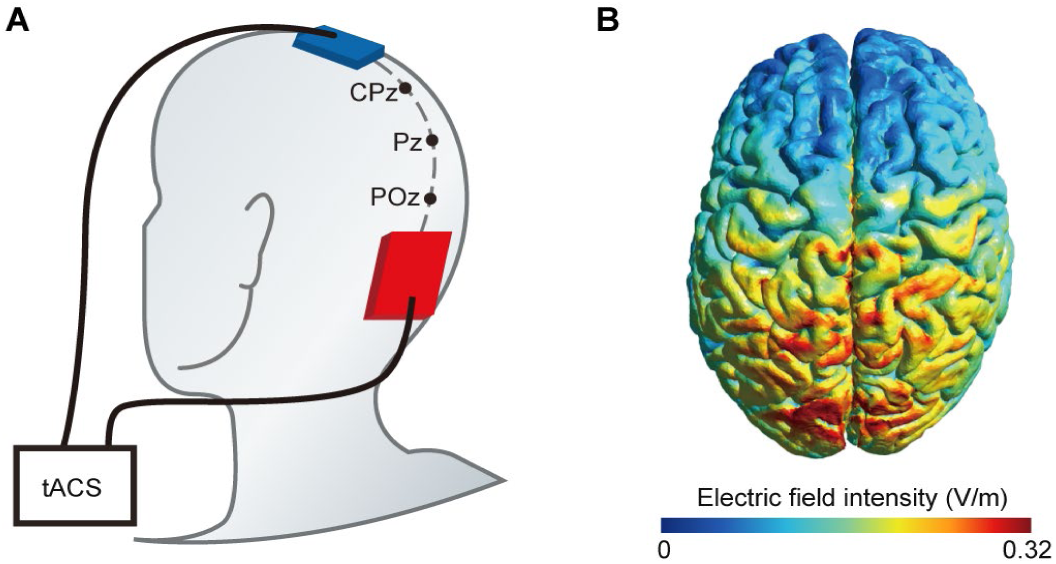
tACS setup. (A) The tACS electrodes were secured to the scalp using elastic bands. The reference electrode (blue) and stimulation electrode (red) were positioned at the vertex (Cz) and posterior occipital area (Oz), respectively. (B) Computational modelling of the induced electric fields was performed using SimNIBS version 4.1.0; montage and anatomical factors influence the resulting electric-field distribution (Opitz et al., 2015). The simulation depicts the spatial distribution and intensity of the electric fields generated during tACS.

**Figure 5—figure supplement 2.**
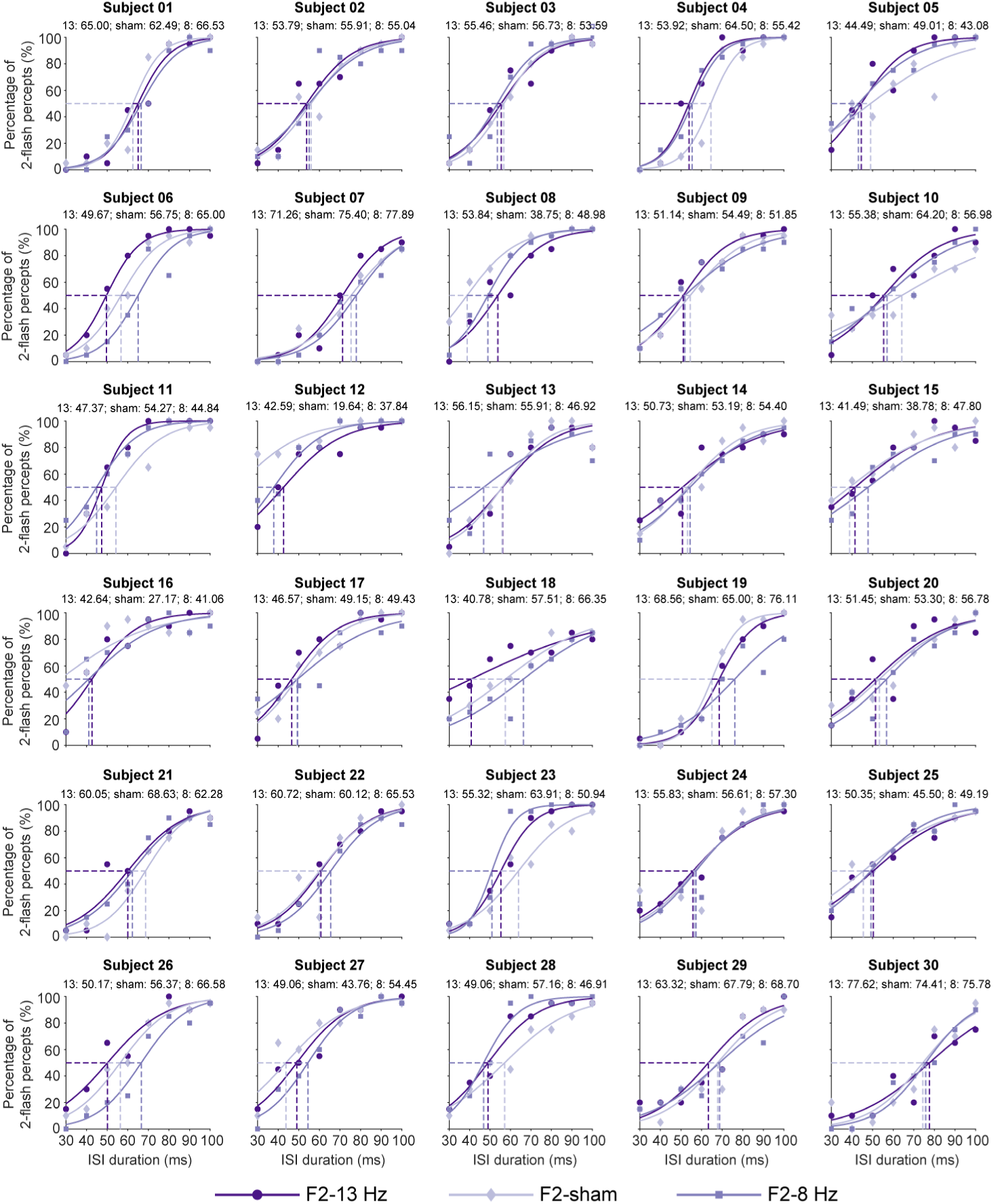
Participant-level psychometric fits in the tACS experiment, F2 condition. Each panel shows one participant’s percentage of two-flash reports as a function of ISI under 13 Hz, sham, and 8 Hz stimulation; symbols are the observed proportions, continuous lines the fitted logistic curves, and vertical 50% threshold markers the fitted fusion thresholds for each session.

**Figure 5—figure supplement 3.**
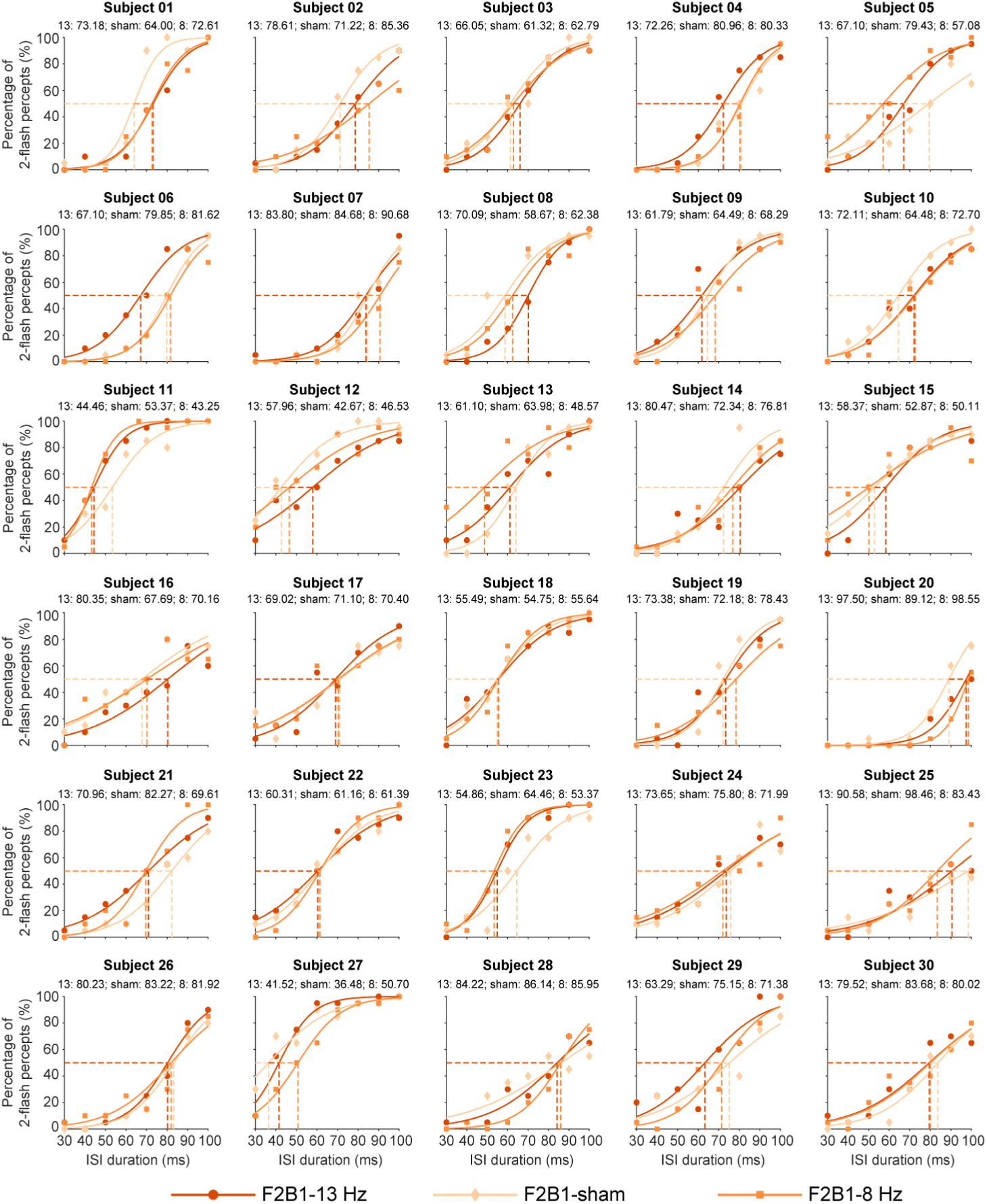
Participant-level psychometric fits in the tACS experiment, F2B1 condition. Each panel shows one participant’s percentage of two-flash reports as a function of ISI under 13 Hz, sham, and 8 Hz stimulation; symbols are the observed proportions, continuous lines the fitted logistic curves, and vertical 50% threshold markers the fitted fusion thresholds for each session.

**Figure 8—figure supplement 1.**
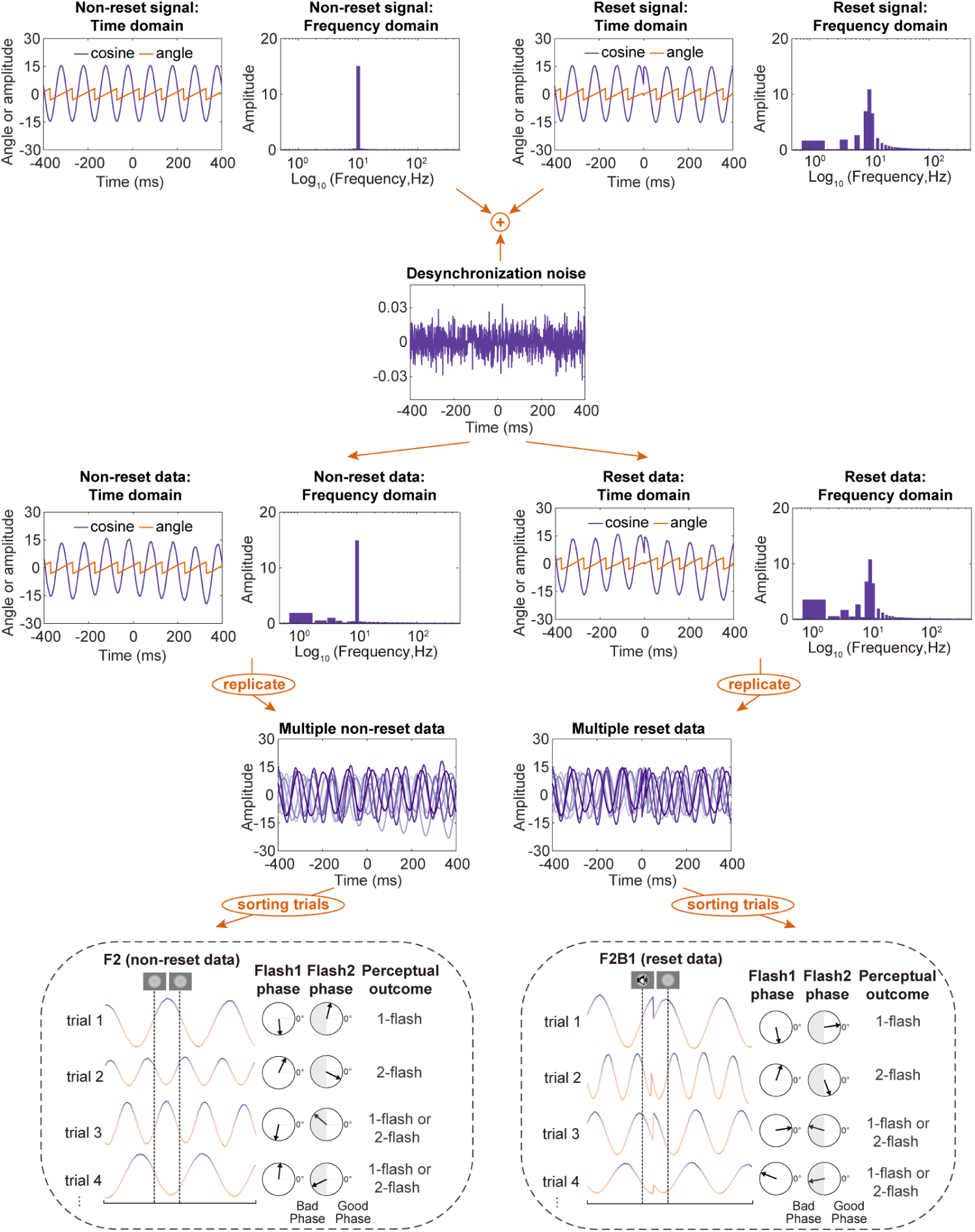
Simulation procedures. Top row: fundamental signals for F2 (non-reset) and F2B1 (reset) conditions in the time and frequency domains; purple lines show the cosine oscillator and orange lines the wrapped phase angle, with the phase reset visible near stimulus onset in the F2B1 signal. Second row: desynchronization noise (the Gaussian innovation parameter e). Third row: addition of the noise to the fundamental signals, giving the final oscillatory data for each condition. Fourth row: repetition of the procedure to generate 1,000 trials for each condition, shown as overlaid traces. Bottom row: assignment of the latent one-flash/two-flash outcome from the cycle relationship between the flashes, followed by phase-dependent report reliability; unfavourable second-flash phases yield a random one- or two-flash report.

**Figure 8—figure supplement 2.**
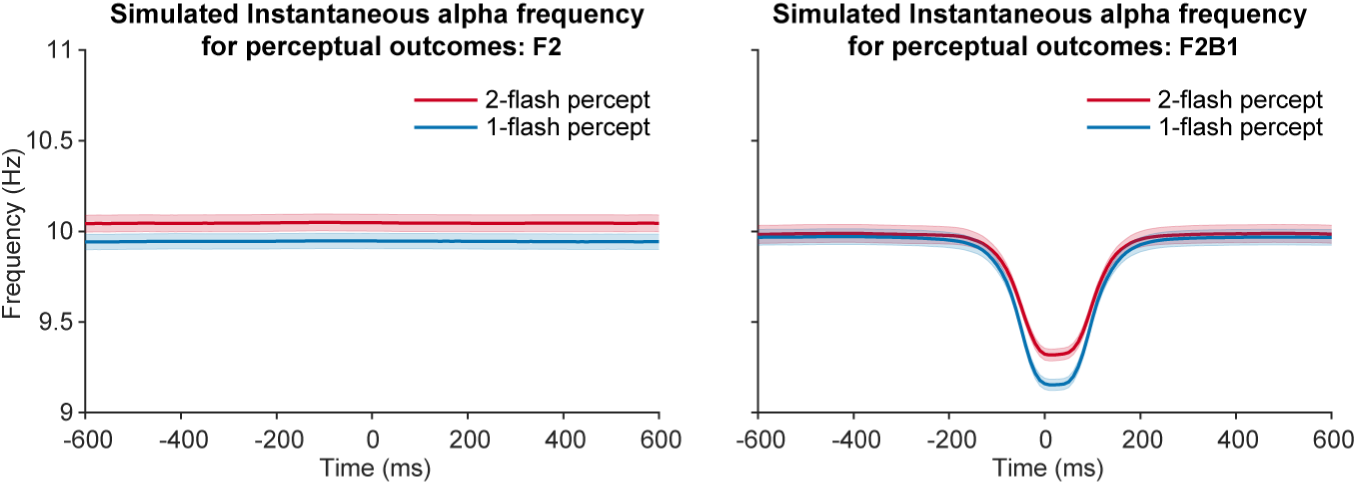
Time-resolved simulated instantaneous alpha frequency for the two perceptual outcomes. Left panel: simulated IAF in F2; right panel: simulated IAF in F2B1. Red and blue lines show two-flash and one-flash percepts, respectively, with shaded areas showing ±1 *SEM* across simulated trials; the modelled decrease near stimulus onset is present in F2B1, and the frequency-percept separation is clearer in F2 than in F2B1.

**Figure 8—figure supplement 3.**
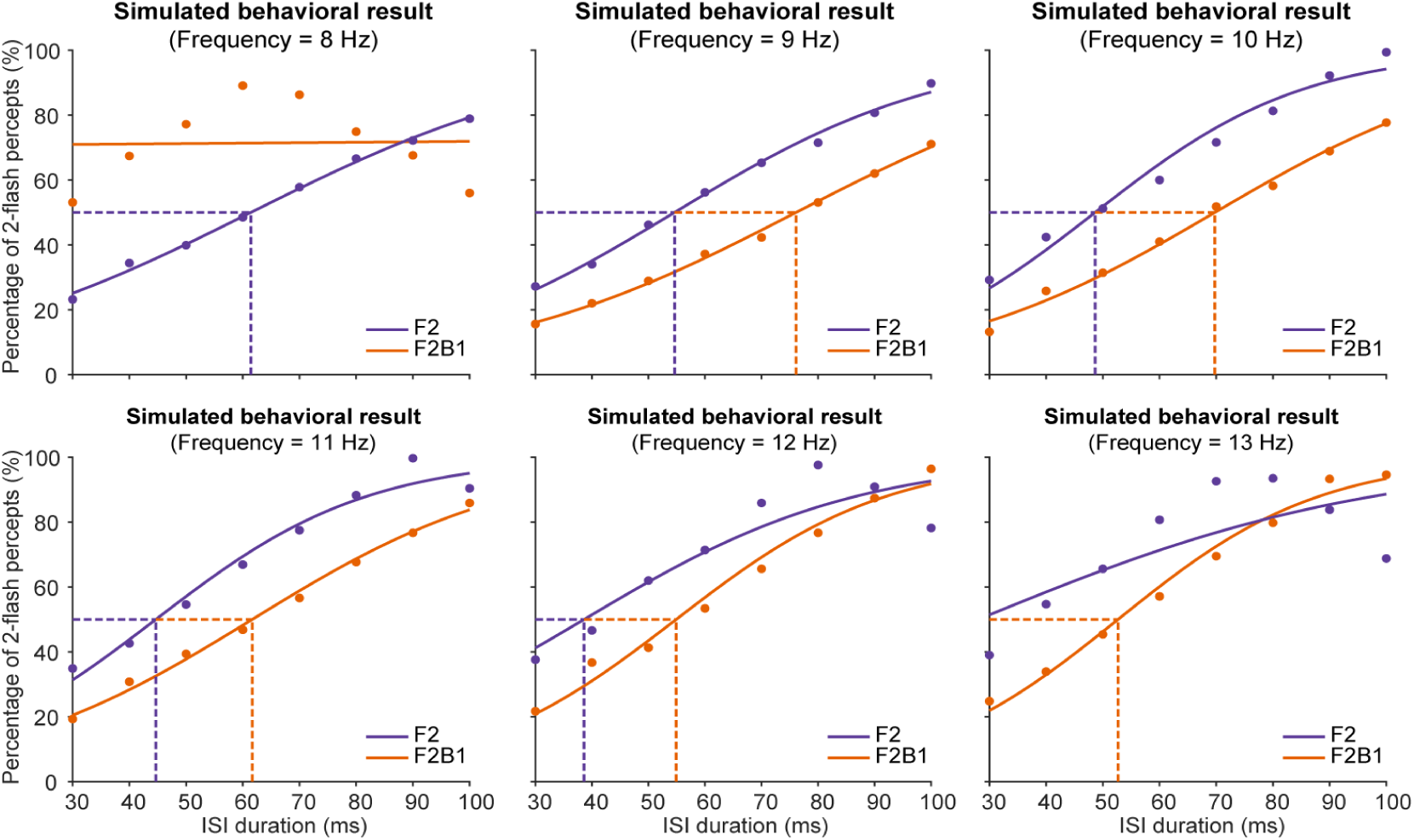
Simulation parameter sensitivity: oscillator frequency. Simulated psychometric curves for F2 (purple) and F2B1 (orange) at fixed oscillator frequencies of 8–13 Hz (25-ms delay); symbols show the simulated percentage of two-flash reports at each ISI and continuous lines the logistic fits. Dashed lines mark the 50% fusion thresholds where the logistic fits were well defined (thresholds were undefined for F2B1 at 8 Hz and for F2 at 13 Hz).

**Figure 8—figure supplement 4.**
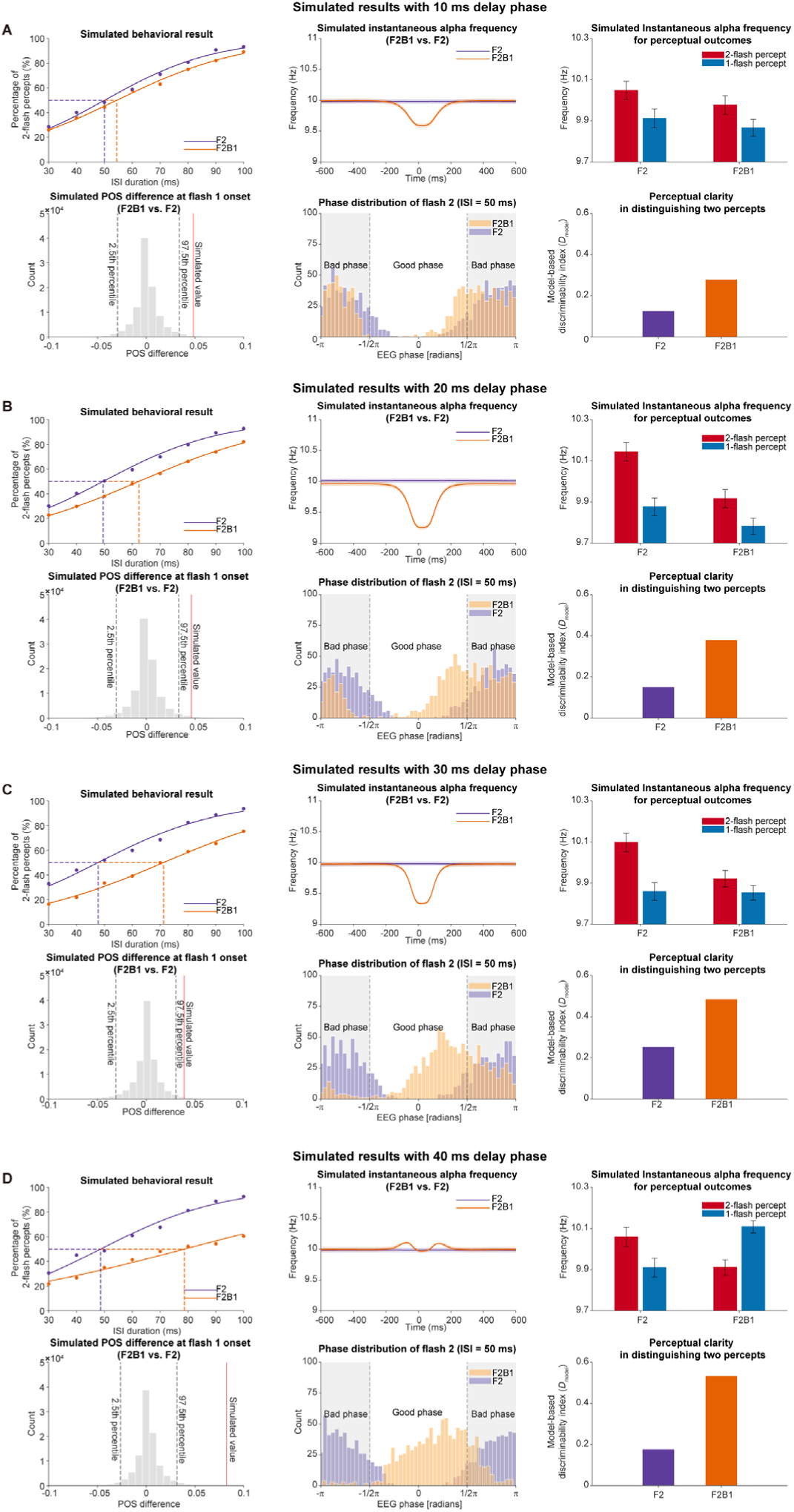
Simulation parameter sensitivity: phase delay. For each delay (10, 20, 30, and 40 ms; panels A–D), the top row shows the simulated psychometric curves (left), the time-resolved IAF for F2 and F2B1 (middle), and the IAF by perceptual outcome (right); the bottom row shows the POS difference at flash-1 onset (left), the phase distribution of the second flash (middle), and the model-based discriminability index (*D*_model_; right). In the psychometric panels, F2 (purple) and F2B1 (orange) are shown with fitted 50% thresholds, and the IAF time courses span −600 to 600 ms. In the POS panels, the grey histograms show the null distributions from 100,000 permutations, the red lines the simulated values, and the dashed black lines the 2.5th and 97.5th percentiles.

**Figure 8—figure supplement 5.**
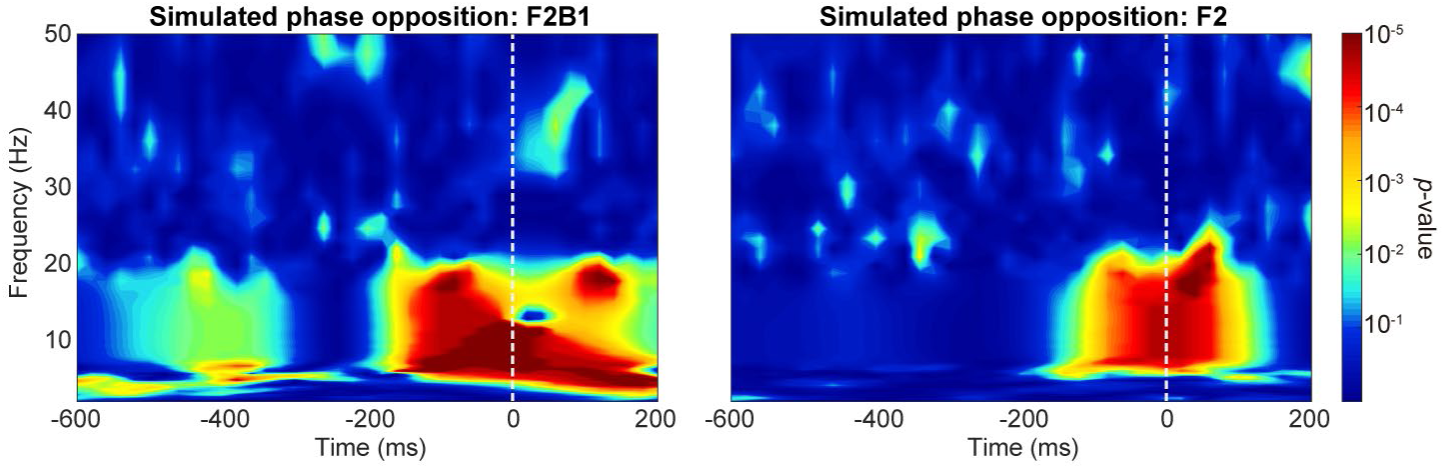
Simulated phase-opposition maps. Left panel: F2B1; right panel: F2. Colour encodes −log10(*p*) from the simulated phase-opposition analysis (shared colour bar on the right), with time on the x axis and frequency on the y axis; both conditions show report-dependent phase separation around stimulus onset, with a larger stimulus-onset difference in F2B1.

